# Neural computation in the brainstem for visceral sensation-driven haemodynamics

**DOI:** 10.1101/2023.08.27.555024

**Authors:** Jiho Lee, Junseung Mun, Sung-Min Park

**Affiliations:** Department of Convergence IT Engineering, Pohang University of Science and Technology (POSTECH), Pohang, 37673, Republic of Korea; Medical Device Innovation Center, Pohang University of Science and Technology (POSTECH), Pohang, 37673, Republic of Korea; Department of Electrical Engineering, Pohang University of Science and Technology (POSTECH), Pohang, 37673, Republic of Korea; Department of Mechanical Engineering, Pohang University of Science and Technology (POSTECH), Pohang, 37673, Republic of Korea; Institute of Convergence Science, Yonsei University, Seoul, 03722, Republic of Korea

## Abstract

The brainstem serves as an intermediary processor of haemodynamic sensations via nucleus tractus solitaries (NTS) in regulating circulatory system. After sensing visceral inputs, the NTS relays information to efferent pathways to modulate peripheral viscera. However, the neural computation mechanism underlying how the NTS processes viscerosensory input remains unknown. Here, we show the computational principles embedded inside the NTS of rats, producing haemodynamic modulation in concert. Our findings demonstrate that the collective dynamics leveraging from neuronal population within the NTS neural circuit encode input-driven haemodynamics. The NTS exhibits the neural trajectory, the dynamical trace of neural states, which is confined to low-dimensional latent space and may represent haemodynamic perturbations. Surprisingly, by normalizing neural trajectory of rats, we found the across-rat common rules for the viscerosensory-information processing by the NTS. Furthermore, the common rules allowed to identify inter-subject variable haemodynamics by quantifying the computational mechanisms in neuro-haemodynamic axis. Our findings provide pioneering insights into understanding the neural computation involved in regulation of visceral functions by the autonomic nervous system.

## Introduction

Haemodynamic homeostasis manages circulatory autonomic outflows and thus regulates the arterial blood pressure (BP) ^1-3^, which is crucial for the survival of mammals, including humans. During haemodynamic regulation, the brainstem is an essential intermediary node between visceral afferent and efferent pathways. In particular, the nucleus tractus solitarius (NTS) is a principal sensory hub that collects vagal sensory cues and promotes the baroreflex which then provides negative feedback for controlling the BP ^4-6^. This emphasizes the computational role of NTS to transform sensory information into the peripheral autonomic tones. However, from the perspective of neural computation, the dynamical principles of the NTS neurons that contribute to haemodynamic regulation remain unknown.

NTS neurons integrate the synaptic inputs from multiple cardiovascular-respiratory afferents ^7-12^. Classical and recent studies have confirmed that the input-driven activation of NTS neurons closely affects changes in BP ^13-17^. However, this single-cellular viewpoint is insufficient for elucidating the influence of NTS on haemodynamics, especially due to heterogeneity in activation patterns across cell types ^18-22^. Then, how do the heterogeneous neurons orchestrate haemodynamics? One possibility is that the collective dynamics of inter-connected network among the NTS neurons may contribute to the haemodynamic perturbation. In this regard, classical studies demonstrated that the NTS is not composed of simple relay neurons between the autonomic sensory and motor peripherals; instead, it is a self-modulating recurrent structure with polysynaptic interconnections ^7,23,24^. This suggests that the input-driven haemodynamics can be governed by the collective dynamics leveraging from the NTS local neural circuit.

Here, we dissect the contribution of neural computation within the NTS to the haemodynamic regulation by investigating the collective dynamics of NTS neuronal population involved in haemodynamic perturbations during viscerosensory input. We demonstrate that the temporally evolving neural trajectory of the NTS resides on a low-dimensional latent space and that the haemodynamic perturbations are encoded in the latent space (Fig. 1a). Building on these theoretical foundations, we propose a biophysically plausible system identification framework for visceral sensation-driven haemodynamics. This framework involves quantifying the neural computation mechanisms with NTS neural circuit model and implementing bi-directional neuro-haemodynamic switching (encoding/decoding) (Fig. 1b). Notably, the bi-directional switching enables the individual model parameter optimization solely using haemodynamic data; thus, obviating the need for invasive neural recordings. Our findings provide insights into computation through neural dynamics within the NTS that underlies haemodynamic homeostasis, which deepens our understanding of the neural bases of autonomic nervous system to regulate peripheral organs.

**Fig. 1.**
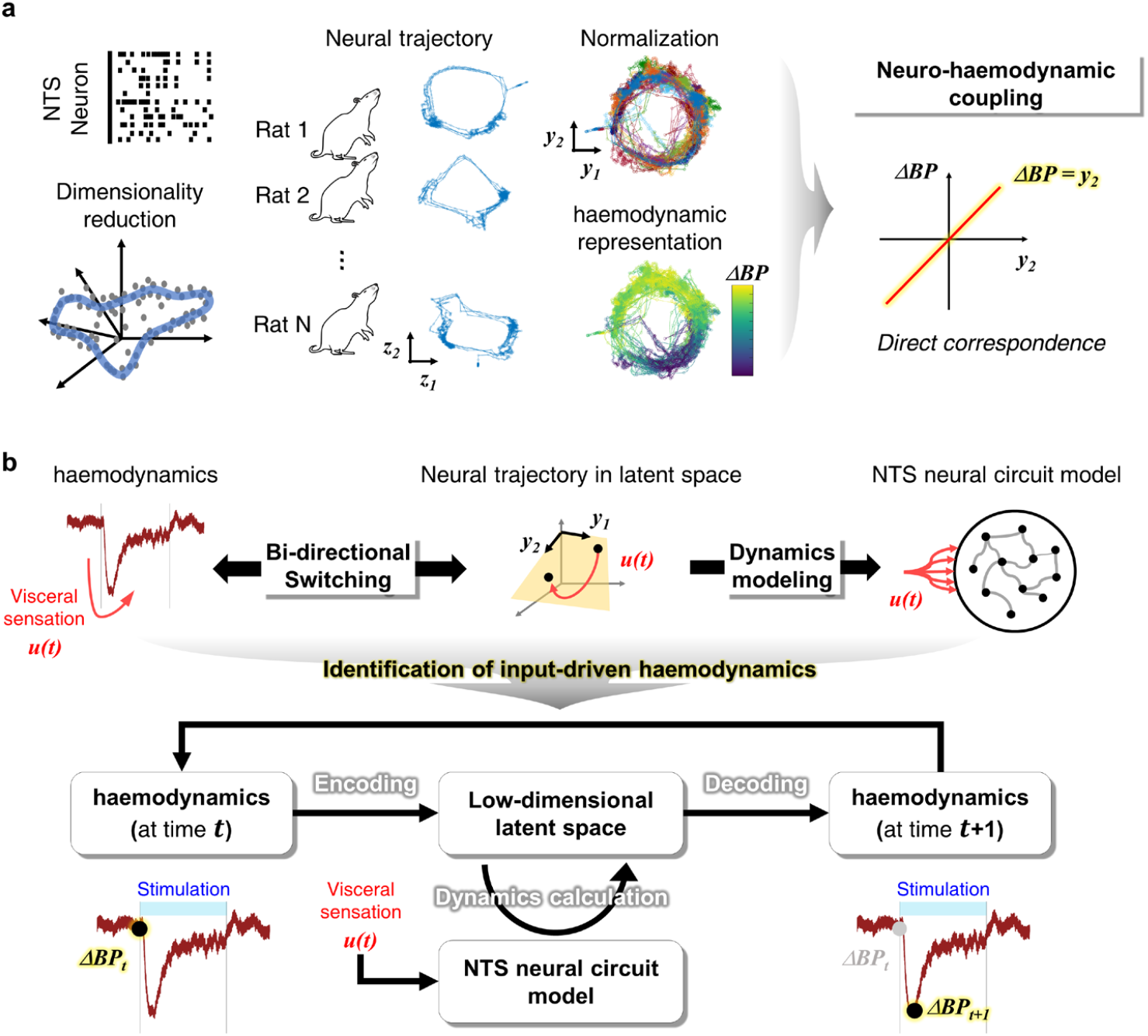
Overview of the study. **a**, Collective dynamics embedded within the nucleus tractus solitarius (NTS) shows ring-shaped neural trajectory residing on the low-dimensional latent space normalized across rats. Vertical axis value of the neural state along the neural trajectory is linearly coupled to viscerosensory input-driven haemodynamic perturbations (representatively ΔBP, change in blood pressure). **b**, Neural computation mechanisms enable to identify a dynamical system underlying the visceral sensation-driven haemodynamics by biophysically modelling the neuro-haemodynamic axis.

## Results

### Latent representation of NTS

We recorded extracellular single-unit activities of NTS and simultaneously measured the femoral arterial BP of the rats (Fig. 2a). Electrical stimuli were applied at solitary tracts projecting to the NTS to imitate viscerosensory inputs, and NTS neurons responsive to the stimulation were selected (*n* = 192; 72% of 266 neurons from 10 rats; Supplementary Fig. 1a-b). The stimulus inputs resulted in temporal evolution of heterogeneous neuronal responses in the NTS (Fig. 2b). The NTS neuronal population comprised 51% activated (*n* = 98), 29% deactivated (*n* = 55), and 20% complex-responding (*n* = 39) neurons of 192 total neurons (Supplementary Fig. 1c). The joint activities among these heterogeneous neurons were examined to determine the effect of inter-neuronal dialogues on heterogeneity across neurons. The pairwise cross-correlations among the measured NTS neurons were higher than that of the dummy data consisting of randomly shuffled time courses of measured neuronal activities. The absolute values (log-scaled) of cross-correlations were as follows: 0.0533 (interquartile range [IQR]: 0.046–0.060) and 0.59 (IQR: 0.48–0.64) for the dummy and measured data, respectively, *p* < 1.83×10^−4^. Shared variances of each neuronal response were highly described with other neurons: 27% (IQR: 22–35%) and 83% (IQR: 76–87%) for the dummy and measured data, respectively, *p* < 1.83×10^−4^ (Fig. 2c). These results indicate that the heterogeneity originates from interconnections formed within the NTS neuronal population, which supports that the NTS accompanies a local neural circuit through the interconnections and corresponds with the finding of previous studies ^18,23^.

**Fig. 2.**
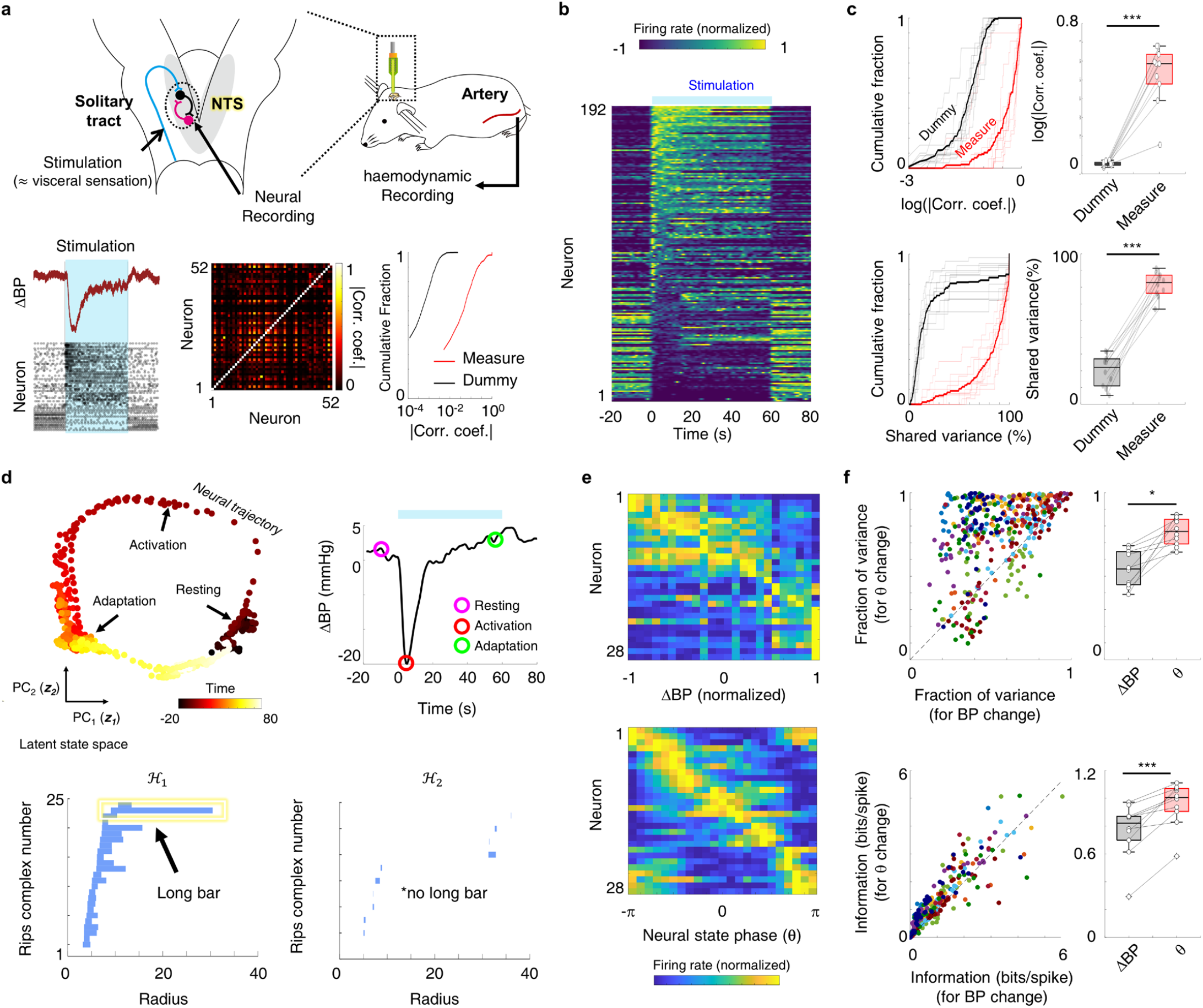
Collective dynamics by the local neural circuit within the nucleus tractus solitarius (NTS). **a**, Illustrations for experimental setup for the neural and haemodynamic recording and their representatives. Neuronal activities among the NTS neurons are cross-correlated compared with those from shuffled dummy data. **b**, Time courses of input-driven NTS neuronal activities. **c**, Statistical analyses for joint activities among the NTS neurons; comparison of pairwise absolute cross-correlation coefficient (top) and shared variance (bottom) between recorded NTS neurons and their shuffled dummy data (*n* = 10 rats; curves: cumulative distributions, light colour: distribution for each rat, and deep colour: mean distribution; box: median ± interquartile range (IQR) and white circles: raw data). **d**, Input-driven neural trajectory from the NTS neuronal population. (Top, left) neural trajectory in low-dimensional latent space (principal component, PC); (top, right) the associated blood pressure (BP) time course; (bottom) Betti barcode. **e**, Tuning curves of single NTS neurons along change in BP (ΔBP) and neural state phase (Δθ). **f**, Capture of single-neuronal activities by ΔBP and Δθ. (Top) fraction of explained variance and (bottom) mutual information (box: median ± IQR). (*: *p* < 0.05 and ***: *p* < 0.001)

To describe how the locally recurrent circuitry within the NTS embeds the collective dynamics converging the joint activities of the neuronal population, we used dimensionality reduction to elicit the neural trajectory in a low-dimensional latent state space that provides compact interpretations and visualizations for the collective dynamics. A key theoretical base of this dimensionality reduction is that the recurrent neural circuit, conveying large information content ^25^, has been extensively established to confine the degree of freedom of the collective dynamics to much lower dimensions than the original number of neurons ^26-31^. After reducing the dimension, we investigated the geometrical topologies and the effective dimension of the neural trajectories in the latent space. We described the topological structure of the trajectories using the persistent homology ^32^ and Betti barcodes visually identified homology groups, each indicating unique topological characteristics (H_0_: point or solid sphere, H_1_: ring, and H_2_: hallow sphere or toroid). As shown in Fig. 2d, we confirmed that the neural trajectory within the NTS neural circuit follows a ring topology, sharply coinciding that the relatively long barcode lifetime in H_1_ (compared with no long bar in H_2_; *z*-scored lifetime of the longest bar: 12.38 [IQR: 10.33–14.01] for H_1_ and 2.50 [IQR: 2.252–3.415] for H_2_; Supplementary Fig. 2a). The effective neural trajectory dimension was specified by investigating how many principal components (PCs) in the latent space were necessary to sufficiently explain the dynamic variance of the trajectory. Our results showed that only the first and second components explained over 86% dynamical variance (mean ± standard deviation(STD) of explained variance: 65.2 ± 8.05% for the first PC and 21.4 ± 6.21% for the second PC; Supplementary Fig. 2b), indicating that the effective latent dimension was bound to two-dimensions (2D). Furthermore, we measured the correlation dimension by plotting the cumulative number of neighbors around a given neural data point while gradually increasing the distance threshold. The increasing order of neighbors versus distance was close to one (mean ± STD of slope of neighbor number versus distance: 0.84 ± 0.031; Supplementary Fig. 2c), corresponding to the ring-shaped topology ^33^.

The NTS neural states along a trajectory in the latent space could discriminate each characteristic activity of single neurons. We compared the tuning curves of neuronal activities along ΔBP and along changes in the neural state, and the results showed that the tuning curve along the neural state distinguished characteristic actions of each neuron (Fig. 2e). Indeed, the neural state phase, an internal variable to quantify the neural state, captured a larger fraction of variance and information involved in single-neuronal activities than did the BP, the external variable (fraction of variance: 0.54 [IQR: 0.45–0.65] and 0.78 [IQR: 0.69–0.84] by the BP and neural state phase, respectively, *p* = 0.0113; mutual information: 0.82 [IQR: 0.70–0.87] and 1.0 [IQR: 0.91–1.1] by the BP and neural state phase, respectively, *p* = 5.83×10^−4^; Fig. 2f). These results indicate that the collective dynamics more successfully describes single neurons than does the haemodynamics in external physical domain.

### Neuro-haemodynamic coupling

In our previous study ^13^, we discovered a surprising and non-intuitive finding that the overall activity of the NTS is correlated with the BP change during stimulation. This direct correlation between NTS activity and visceral output suggested that the computation by the NTS could determine the temporal evolution of input-driven haemodynamics. This hypothesis introduced a new viewpoint on the computational role of NTS to modulate circulatory system, and thus we examined how the NTS encodes input-driven haemodynamics.

To elucidate the neuro-haemodynamic coupling protocol, we investigated how precisely single-neuronal activities predict the BP time course. We found that single NTS neurons did not predict changes in BP (ΔBP) (Fig. 3a-b). Then, we decoded the neural trajectory in latent space into haemodynamics on the basis of recent studies that suggested an emerging approach of ‘computation-through-dynamics’ ^31^. This approach has provided a good understanding of how the brain encodes body functions related to cognitions and behaviors, including the head directions of rodents encoded along the ring structure in the head-direction cells ^32^, spatial recognition encoded along the toroidal structure in the grid cells ^34^, and motor reactions cued by a three-dimensional neural trajectory in the cortex ^35^. Analogously, we assumed that the autonomic body functions, especially haemodynamics in this study, could be encoded by the neural trajectory in latent space as in cognitive and motor functions. We disentangled the relationship between the latent space of the NTS and the haemodynamic perturbations, finding that the latent space was linearly coupled to the ΔBP (Fig. 3c). Each neural state along the ring-shaped neural trajectory did not represent a unique value of BP. Rather, the ΔBP was represented in the latent space following a linear subspace that we refer to as ‘decoding space’ (see ‘Disentanglement of neuro-haemodynamic axis’ in Method session). Neural decoding through the linear protocol showed high prediction accuracy for the BP time course (89%; Fig. 3d) and a lower mean squared error than did single neurons (log scale of mean squared error: -0.799 and -1.96 for single neurons and neural decoding, respectively; Fig. 3e). These indicate that the NTS encodes ΔBP using collective dynamics. Moreover, our statistical analyses support the evidence by presenting a much higher prediction accuracy using neural decoding than using single neurons (prediction accuracy: 0.28 [IQR: 0.24–0.31] and 0.78 [IQR: 0.70–0.93] for single neurons and neural decoding, respectively, *p* < 1.83×10^−4^; Fig. 3f) and a smaller prediction error (log scale of mean squared error: -0.75 [IQR: -0.84 to -0.62] and -1.22 [IQR: -1.78 to -1.01] for single neurons and neural decoding, respectively, *p* = 2.46×10^−4^; Fig. 3g).

**Fig. 3.**
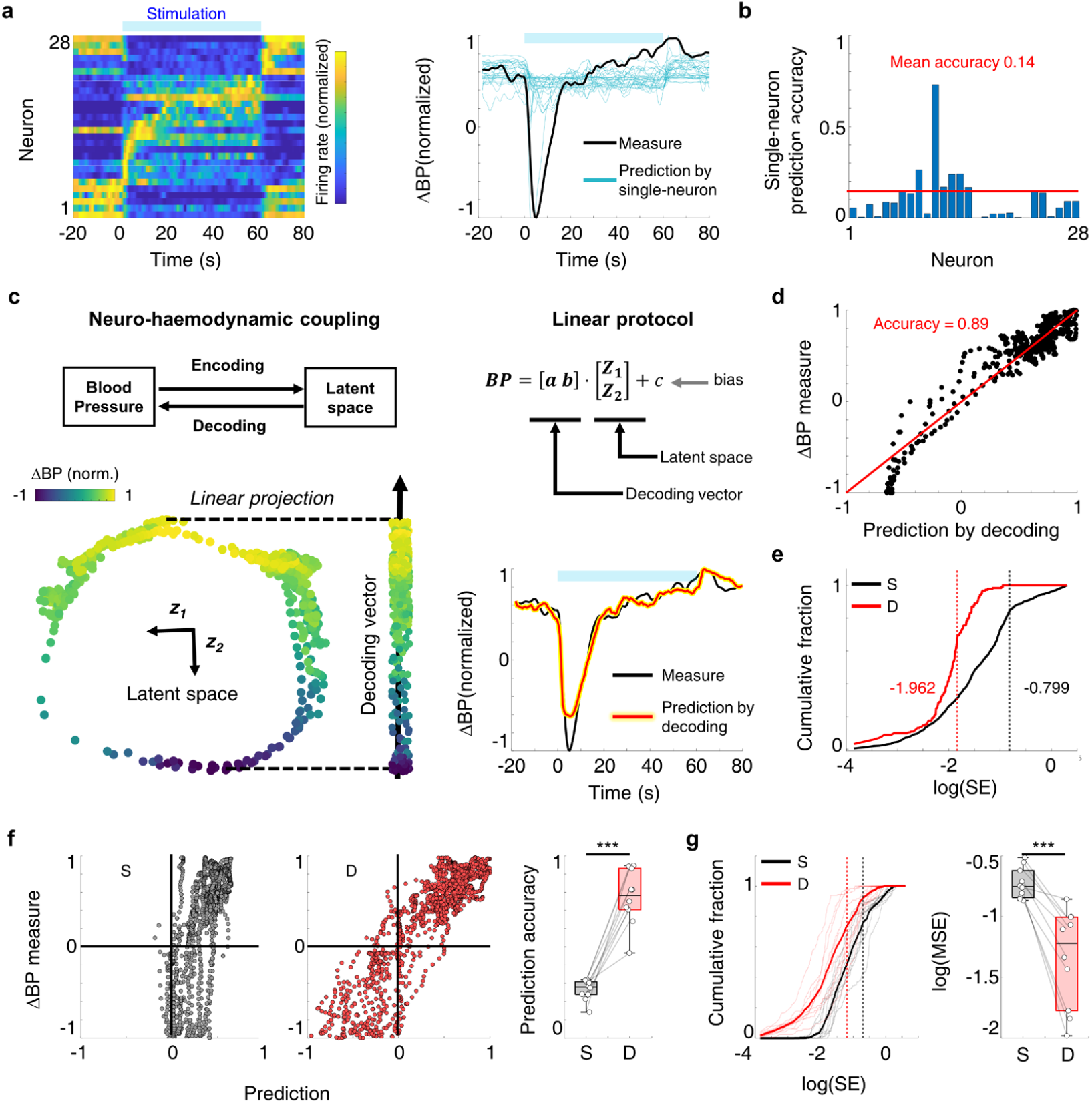
Relationship between nucleus tractus solitarius (NTS) and input-driven haemodynamics. **a**, Prediction of blood pressure (BP) time course by each single-neuronal activity. (Left) single-neuronal activities; (right) prediction and measured BP time course. **b**, Prediction accuracy with single neurons (bar: prediction accuracy for each neuron from rat 1). **c**, Neuro-haemodynamic coupling between NTS latent space and input-driven haemodynamics. (Left) representative illustration of the coupling; (right) prediction of BP time course with neural decoding through the latent space. **d**, Prediction by the neural decoding for the measured BP time course (black dots: data point pair for each time step; red line: linear regression). **e**, Cumulative distribution of squared error (SE) of prediction by single neurons (S) and prediction by neural decoding **d**, (vertical dotted line: mean of error distribution). (**f-g**). Statistical analyzes for prediction accuracy to compare predictions by single neurons and by neural decoding (*n* = 10 rats; **f**, prediction accuracy and box: median ± IQR; **g**, prediction error described by SE for each rat and box: median ± IQR for mean squared error [MSE]). (***: *p* < 0.001)

Next, we investigated the input-driven characteristics of various haemodynamic functions with consideration of the anatomically known efferent pathways downstream ^2^ from the NTS to haemodynamic peripherals (Supplementary Fig. 3a). Indeed, the change in BP evolved in delay following stimulation onset (mean±STD of delay: 4.10±2.22 second; Supplementary Fig. 3b), designating the propagation time from the NTS to peripherals. We observed input-driven haemodynamics in several functions, including the spectral power of heart rate variability which indicates autonomic tones (low and high frequency power for sympathetic and parasympathetic tone, respectively), heart rate, and BP (Supplementary Fig. 3c-d). Thereafter, explanations for perturbations in the haemodynamic functions by single-neuronal activities and collective dynamics were analysed. The single-neuronal activities did not capture any haemodynamic perturbations, whereas the neural decoding could capture the heart rate with high prediction accuracy, similar to that of BP prediction (Supplementary Fig. 3e-f). These suggest the determination of peripheral modulation even at the level of the NTS rather than other pre-automatic nodes such as rostral ventrolateral medulla and dorsal motor nucleus of the vagus after the NTS^2^.

### Across-rat common rules for NTS

Despite the commonalities of ring-shape topology and linear neuro-haemodynamic relationship as shown in Fig. 3, each rat displayed a distinct neural trajectory, reflecting heterogeneities in the collective dynamics and the decoding vector (Fig. 4). Specifically, geometric heterogeneity encompassed variations in the positions of center points and the scales of axes (Supplementary Fig. 4). The dynamic heterogeneity encompassed variations in the positions of stable points of neural state (before and after stimulation for resting and recovering, respectively) and the rotational directions across rats (during stimulating), as represented by time courses of the neural state phase. Angles of the decoding vectors were randomly distributed, which is consistent with the difference in the decoding vector (mean ± STD of the angle of decoding vectors: 1.38π ± 0.28π rad; Supplementary Fig. 4c).

**Fig. 4.**
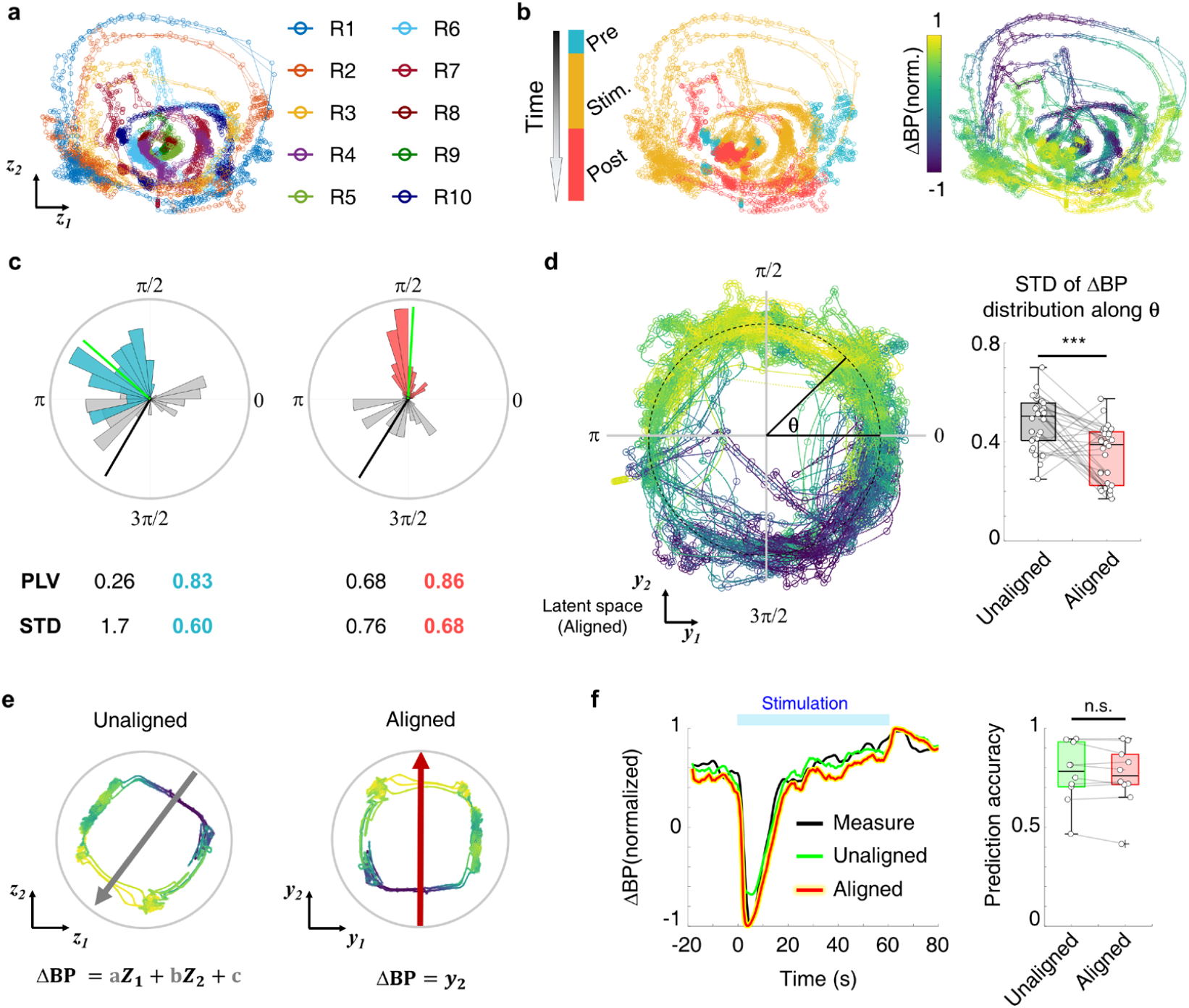
Normalization of different neural trajectories from the nucleus tractus solitarius (NTS) across rats. **a**, Rat-specific neural trajectories (*n* = 10 rats). **b**, Heterogeneity in dynamics of neural states (left) and neuro-haemodynamic coupling (right; ΔBP: change in blood pressure). (**c-d**) Across-rat normalization of neural trajectory through alignment. **c**, Polar histogram of distribution of neural state phase (*n* = 30 bins; unaligned distribution: grey; aligned distribution: cyan during pre-stimulation and peach during post-stimulation; mean of distribution: black line for the unaligned and green line for the aligned). **d**, (Left) Distribution of haemodynamic representation in the aligned latent space; (right) statistical analyses (*n* = 30 bins) for standard deviation (STD) of ΔBP along the neural state phase (θ) (box: median ± IQR and white circles: raw data). **e**, Illustrations for the common protocol of neural decoding enabled by alignment (*z*_*1*_ and *z*_*2*_: unaligned latent space; ***y***_***1***_ and ***y***_***2***_: aligned latent space; grey coloured a, b, and c: regression coefficients for rat-specific neural decoding). **f**, Comparison between neural decoding of rat-specific unaligned latent space and across-rat aligned latent space (*n* = 10 rats; box: median ± IQR and white circles: raw data). (***: *p* < 0.001 and n.s: not significant).

Interestingly, we found that the rat-specific neural trajectories and their haemodynamic representations can be normalized across rats. Each rat-specific neural trajectory in latent space (*z*_1_, *z*_2_) was normalized by aligning onto the unit circle in a latent space (*y*_1_, *y*_2_; Supplementary Fig. 4f). Changes in the neural state phase in the aligned latent space showed low variability of input-driven dynamics across rats. The alignment increased phase-locking values (PLV) of the distribution of phase during the resting and the recovering (resting: from 0.26 to 0.83; recovering: from 0.68 to 0.86) and decreased standard deviation (STD) (resting: from 1.7 to 0.6l; recovering: from 0.76 to 0.68; Fig. 4c). However, despite the low inter-subject variability, we also found that the dynamics were not identical across rats (Supplementary Fig. 4g); this was consistent with the heterogeneous haemodynamic responses of rats even with same input application (Supplementary Fig. 3c). As for the neuro-haemodynamic coupling, the representations of change in BP along the aligned neural trajectories significantly lowered STD across rats than those of the unaligned trajectories (n = 30 bins, STD distribution: 0.52 [IQR: 0.40–0.57] and 0.37 [IQR: 0.24–0.45] for unaligned and aligned trajectories, respectively, *p* = 0.00258; Fig. 4d). Moreover, we evaluated the performance of input-driven BP prediction with the decoding of the aligned latent space, compared to the decoding of rat-specific unaligned latent space. We chose the across-rat common decoding space from the aligned latent space as the vertical axis for the simplicity of calculation (Fig. 4e). There was no significant difference in BP prediction accuracy of decoding between the unaligned (non-normalized) and the aligned (normalized) latent spaces (*p* = 0.232; Fig. 4f), indicating that inter-individually variable haemodynamics were sufficiently captured by the common protocol of neuro-haemodynamic coupling using the pre-determined normalized latent space.

Next, we investigated the principles underlying inter-rat heterogeneities in the neural trajectories and the degree of coupling (strong or weak; quantified with the aforementioned prediction accuracy) and suggested that although an identical NTS neural circuit is shared across rats, the difference in recorded subpopulations for each rat, known as ‘recording instability’ ^36^, introduces the heterogeneity. We first showed that when subpopulations were extracted from a single neuronal population through random sampling, these subpopulations exhibited distinct neural trajectories. The differences encompassed the geometric distortion and the various BP prediction accuracies (*n* = 100 random subpopulations; Supplementary Fig. 5). Then, we demonstrated that the heterogeneous neural trajectories can be aligned onto the unit circle while not changing their own prediction accuracy for each subpopulation (Supplementary Fig. 5e). Considering the similarity between the rat-and subpopulation-specific trajectories, we deemed it plausible that the heterogeneous neural trajectories may not originate from inter-rat variable characteristics of NTS neural circuit, but rather arise from different subpopulations within the across-rat shared NTS neural circuit. Furthermore, we calculated an overall neural trajectory by gathering all recorded neurons from different rats and this also followed the 2D ring-shape as for each rat trajectory (Supplementary Fig. 6a-d). The conservation of ring topologies in both overall population and subpopulations is consistent with the previously-mentioned relationship between the original population and its subpopulations. We also showed that the neural trajectory of one rat could be replicated by subsampling the overall neuronal population even not including the neurons recorded from the rat (Supplementary Fig. 6e), indicating that common neurons are recorded in different rats. These results support our hypotheses that the NTS neural circuit is common across rats and the differences in dynamical and functional characteristics depend on recording quality. In other words, we suggest three significant across-rat common rules for NTS: 1) viscerosensory input-driven collective dynamics follows ring-shaped neural trajectory; 2) input-driven haemodynamics is sufficiently captured by neural trajectory on across-rat normalized latent space; 3) properties of NTS neural circuit are similar across rats.

### System identification of input-driven haemodynamics

Our findings regarding the neural computations in the NTS enabled us to identify the input–output dynamic system underlying the haemodynamic perturbation (output) after NTS receives visceral sensation (input), the process called as ‘system identification’ ^37^. Specifically, we formulated a biophysically-plausible framework for computational dynamics that infers the input-driven haemodynamics by quantitatively modelling the dynamics in NTS neural circuit and neuro-haemodynamic coupling, which we named as the haemodynamics model based on barosensory input-driven neural dynamics (H-BIND; Fig. 1b and 5a). The workflow of H-BIND starts with transforming a current haemodynamic value (**BP**_*t*_) into a state in latent space 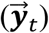 (process of ‘Encoding’). Then, the H-BIND updates the next latent state 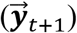 using the NTS neural circuit model (process of ‘Input-driven dynamics calculation’). Finally, the H-BIND returns the updated haemodynamic value (**BP**_*t+1*_) by inverse-transforming the updated latent state (process of ‘Decoding’). Thus, the system identification process for the H-BIND was subdivided to build the NTS neural circuit model and the neuro-haemodynamic encoder/decoder.

**Fig. 5.**
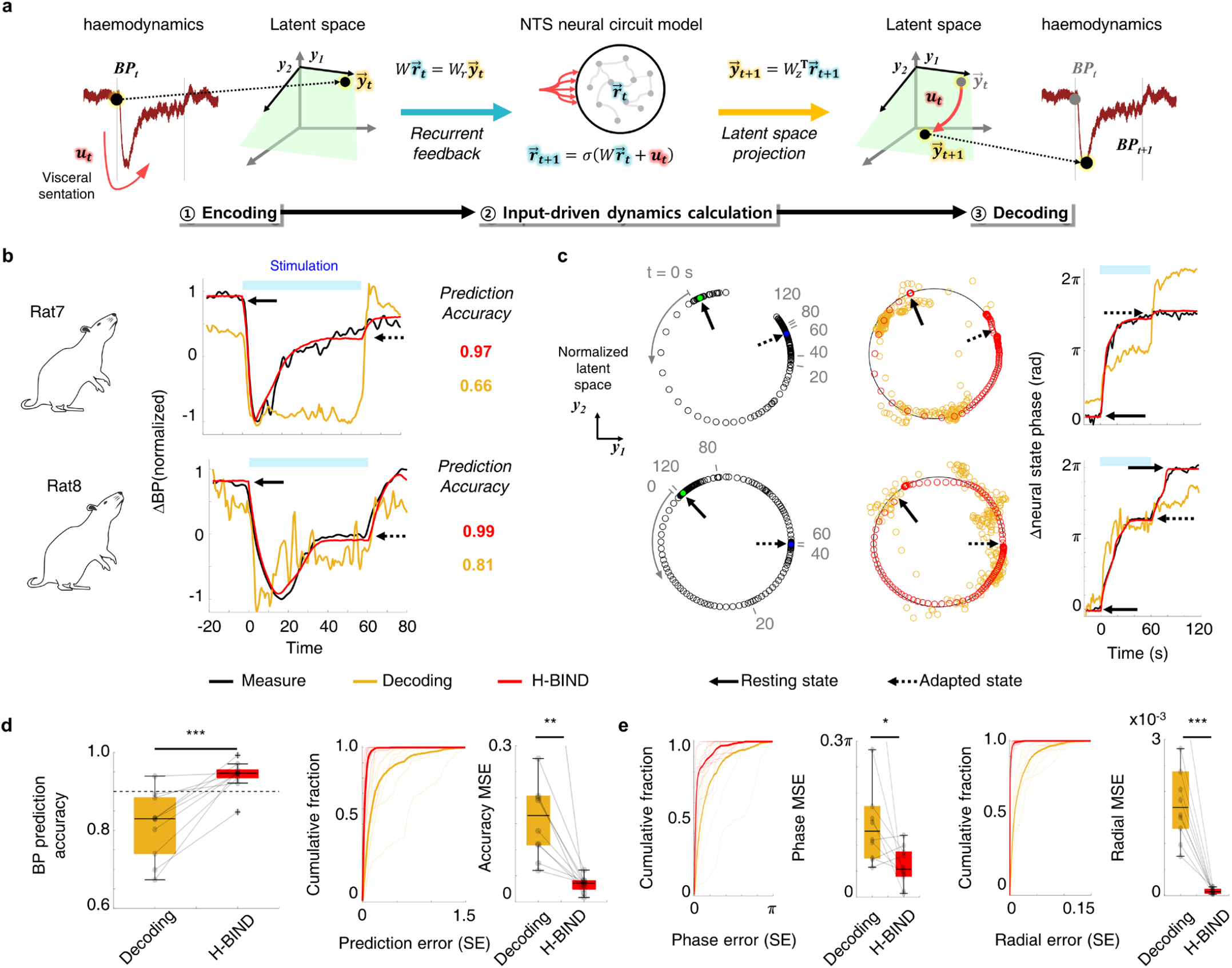
System identification of input-driven haemodynamics using neural computation mechanisms in the nucleus tractus soliatrius (NTS). **a**, Illustration for the concept of the haemodynamics model base on barosensory input-driven neural dynamics (H-BIND). **b**, Prediction by H-BIND for inter-individually variable temporal dynamics of blood pressure (BP) compared with that of empirical neural decoding from the NTS recordings. **c**, Comparison of estimation for neural trajectory in across-rat normalized latent space using H-BIND with that of empirical neural decoding; ground-truth neural trajectory is reconstructed from BP time course using bi-directional neuro-haemodynamic transformation. (**d-e**). Statistical analyses for the haemodynamics prediction. **d**, Prediction accuracy and error for change in blood pressure (ΔBP; *n* = 10 rats; box: median ± IQR and white circles: raw data; prediction error described by cumulative distribution of squared error (SE) for each rat and box: median ± IQR for mean squared error (MSE)). **e**, Prediction errors (radial and phasic error) for neural trajectory in normalized latent space (*n* = 10 rats; errors described by cumulative distribution of SE for each rat and box: median ± IQR for MSE). (*: *p* < 0.05, **: *p* < 0.01, and ***: *p* < 0.001)

The NTS neural circuit was modelled with a recurrent neural network (RNN) ^38^. We found that multivariate linear regression of NTS singe-neuronal activities sufficiently expressed the change in latent space (Supplementary Fig. 7a-b), enabling the design condition for the RNN that the activities of each unit, equal to neuron in real NTS, linearly exhibit the neural trajectory in latent space (Supplementary Fig. 7c). The trained RNN successfully modelled the dynamics of the real NTS neural circuit, replicating the measured single-neuronal activities and neural trajectory (Supplementary Fig. 7d-g). As for the neuro-haemodynamic encoder/decoder, we established bi-directional switching between the haemodynamic perturbation and neural trajectory through the pre-determined across-rat normalized latent space. The aforementioned across-rat rule that the normalized latent space sufficiently suggests the bi-directionality of neuro-haemodynamic coupling, implying that the reversal procedure from haemodynamics into normalized latent space is possible. Furthermore, this allowed the proposed H-BIND framework to be individually optimized by training the RNN with the reconstructed neural trajectory from recordings of haemodynamic parameters such as BP, thus facilitating the advantages of not requiring severely invasive NTS neural recordings (Supplementary Fig. 8b-c).

We evaluated the performance of H-BIND to predict different input-driven haemodynamics across rats (Fig. 5b). In two different rats, the H-BINDs individually optimized for each rat successfully predicted the temporal evolutions of haemodynamic perturbation. Compared with the empirical estimation by neural decoding based on the measured NTS activities, the prediction accuracy was much higher for H-BIND. The neural trajectories encoded from the haemodynamic recordings were also more accurately predicted by H-BIND than by neural decoding (Fig. 5c). Our statistical analyses support the superior performance of H-BIND for haemodynamic prediction (prediction accuracy: 0.83 [IQR: 0.74–0.88] and 0.95 [IQR: 0.93–0.96] for empirical neural decoding and H-BIND, respectively, *p* = 0.00195; prediction error: 0.17 [IQR: 0.11– 0.20] and 0.036 [IQR: 0.024–0.041] for empirical neural decoding and H-BIND, respectively, *p* = 0.00195; Fig. 5d) and neural trajectory prediction (phase error: 0.13 rad [IQR: 0.075–0.17] and 0.054 rad [IQR: 0.040– 0.087] for empirical neural decoding and H-BIND, respectively, *p* = 0.0371; radial error: 0.017 [IQR: 0.013– 0.024] and 0.00072 [IQR: 0.00041–0.0012] for empirical neural decoding and H-BIND, respectively, *p* = 0.00195; Fig. 5d-e).

## Discussion

This study elucidates the mechanisms of neural computation within the NTS processing viscerosensory information that underlies the regulation of blood circulation through the autonomic nervous system. The joint interactions of heterogeneous NTS neurons unfold the collective dynamic neural trajectory in low-dimensional latent space. We investigated the characteristics of the NTS neural trajectory, including the ring-shaped geometry spanning on 2D latent space that encodes haemodynamic perturbations; thus, demonstrating the neuro-haemodynamic entanglement through the latent space leveraging from the collective dynamics. Our findings provide pioneering insights into the computational principles within the NTS. The direct NTS-haemodynamic dialogue indicates a close association between peripheral viscera and the NTS, bypassing other intermediate efferent nodes. While the NTS has long been considered as a simple relay for the sensory information into pre-autonomic nodes ^8,39,40^, the early involvement of the NTS in determining haemodynamics suggests its governing role in balancing autonomic tones within the circulatory system. Additionally, previous studies have reported the significant role of the NTS in regulating other autonomic functions such as feeding ^*41*^ and respiration ^*42*^, which aligns with our perspective.

We emphasize a significant viewpoint for the neural computation embedded in the autonomic nervous system; specifically, the similarity to that in other brain hubs such as the cortex and the hippocampus that administrate motor ^33,35^ and cognitive functions ^34,43^. This improves our comprehensive understanding about the brainwide usage of neural computations through collective dynamics for brain-peripheral interfaces. In this context, preliminary proceedings in other brain hubs suggests future works that may lead us toward deeper understanding of unknown computational mechanisms underlying physiological functions controlled by the autonomic nervous system. For example, the plausible explanation for unknown type of dynamics that the NTS neural trajectory follows can be an ‘attractor’ leveraging from recurrent circuitry as shown in hippocampus ^31^. We may need to consider long-term monitoring over days to observe slowly-evolving characteristics of the NTS neural trajectory as in motor cortex ^43^ and hypothalamus ^44^. Another significant inquiry is the context-dependent collective dynamics ^30^ unfolded with multi-type visceral sensations and mental conditions. The possibility of contextual-dependent effects is supported by classical investigations demonstrating that NTS neurons integrate multiple input sources such as cardiorespiratory ^11^ and stress-hormonal inputs ^45^. Additionally, although we targeted the circulatory system owing to abundant evidence of NTS-haemodynamic coupling, the existence of inter-neuronal connections spread overall the NTS, as reported in previous studies ^9,18,23^, provides motifs toward large unknowns concerning NTS contributions to other autonomic functions such as in the gastrointestinal and laryngeal systems. Thus, substantial future works are required to discover sharp profiles of the collective dynamics within the NTS regulating autonomous system.

We found the across-rat common rules in the neural computation mechanisms of the NTS underlying the temporal evolution of the viscerosensory input-driven haemodynamics. Aligning neural trajectories onto pre-determined normalized latent space across rats unveil that the normalized latent space sufficiently captures haemodynamic perturbations for each rat; indicating the commonalities in the shape of collective dynamics represented by neural trajectory and the neuro-haemodynamic coupling protocol even implying bi-directional switching. We emphasize that this calibration for the neuro-haemodynamic dialogue promotes a biophysically-plausible and individualized system identification framework (H-BIND). Specifically, the common rule for collective dynamics enabled to construct a generalized model architecture for the NTS neural circuit. Notably, the model parameters could be individually optimized using an inter-individually variable neural trajectory that was obtained by the bi-directional switching of haemodynamic perturbations; indicating no need of invasive direct neural recording. By capturing the intrinsic dynamics of the neuro-haemodynamic axis, H-BIND elicits remarkable predictive ability for haemodynamic responses to visceral sensation compared with that of empirical neural decoding. This fosters a perspective that H-BIND more accurately estimates the input-driven dynamics in the neuro-haemodynamic axis than decoding measured neural data; potentially due to stable reconstruction of collective dynamics based on across-rat common rules whereas empirical decoding is restricted by recording instability (Supplementary Fig. 5).

Although the limitation on the current stimulus parameters requires further study to optimize H-BIND for variable parameters such as stimulus intensity and frequency ^10^, H-BIND poses the potential to improve our understanding of neural computations involved in autonomic regulations. The accurate neuro-haemodynamic estimation offered by H-BIND could aid in uncovering putative computational mechanisms from the single cell to population levels, even in subtle perturbations ^46^. Additionally, the proposed H-BIND can also have significant implication into the medical technologies, specifically for neuromodulation therapies for circulatory disorders such as hypertension ^47^. The individually-optimizable H-BIND framework could serve as a personalized testbed, allowing the examination of stimulation effects in accordance with patient-dependent conditions. Overall, we envision H-BIND as a potential bridge connecting our understanding of neural computation mechanisms with their extensive future medical applications.

## Methods

### Animals

All surgical and experimental protocols were approved by the Institutional Animal Care and Use Committee at Pohang University of Science and Technology (POSTECH-IACUC; approval number: POSTECH-2021-0017 and POSTECH-2022-0020) and followed the animal experiment guidelines from the NIH Guide for the Car and Use of Laboratory Animals. We used the adult male Sprague-Dawley rats (age: 8-10 weeks; weight: 270-320 g; provided by Orient Bio, Rep. of Korea, colony reference at Charles River Laboratories, and by Koatech, Rep. of Korea, colony reference at Envigo). We performed surgeries with the rat subjects for non-survival acute experiments, and thus at the end of experiments, the subjects were euthanized using CO_2_ gas.

### Surgery

Before the surgical procedures, the rat subjects were anesthetized with intraperitoneal injection of customized urethane solution (injection does: 1.2-1.4 g/kg; density: 0.2 g/mL diluted in saline solution; urethane: purchased from Merck (Sigma Aldrich), product code U2500) after initial induction using isoflurane (5% in O_2_ flow rat 0.4-0.6 L/min; 5-10 min). Supplemental doses of 0.3-0.6 g/kg were additionally injected if needed. After anaesthesia, left femoral artery was cannulated with polyethylene tube (SP-10, Natsume Seisakusho Co.) for arterial BP recording. The subjects underwent the surgeries to expose the brainstem as described in previous studies^22,48,49^. Tracheal intubation was administrated only if the respiration of subject was not stable. The subjects were placed on stereotaxic frame (68002, RWD Life Science), and the craniotomy was performed to plant the reference and ground anchor screws for electrophysiological recording. Thereafter, the placement of subject was changed to permit sharp flexion of the neck for a surgery to approach the dorsal surface of medulla in brainstem. Dorsal neck muscles were incised and retracted to expose the atlanto-occipital membrane. The membrane was incised and part of occipital bone was removed to clearly expose the rhomboid fossa.

### Solitary tract stimulation

Our stimulation protocol followed the method described in our previous study^48^. In brief, we targeted the solitary tracts into the NTS^50^ to simulate the barosensory input into the NTS. The stimuli were delivered with an arbitrary waveform generator (IZ2-32, Tucker-Davis Technologies (TDT)), which is electrically isolated by using a battery (LM48M-250, TDT). We used the customized bipolar electrode fabricated with two enamel-encapsulated stainless still wires (diameter: 150 um, GoodFellow). The electrode was fixed at the stereotaxic arm with a supporter structure made in custom. Stimulation used the bipolar square waveform (pulse width: 100 us; frequency: 20 Hz; intensity: 150-250 uA, differently chosen along individual subjects as driving haemodynamic responses such as the decrease of blood pressure). Although the stimulus frequency is significant factor to determine the properties in temporal dynamics of NTS neural activity^51,52^ and even haemodynamics, especially the blood pressure^15^, we chose the 20 Hz, representing complex dynamic patterns; for an example, the blood pressure first sharply decreases, slowly recover up to pre-stimulation baseline, and finally adapts to the stimulation. Stimulation was applied for 60 s to evoke sufficient dynamic perturbations both in the NTS neural activities and haemodynamic functions. We ensured for 10 min before stimulation experiments to stabilize and thus prevent unknown fluctuations in neuro-haemodynamic signals.

### Electrophysiological neural recording

After exposing surgery of dorsal surface of brainstem, we recorded the NTS neural activities following the methods shown in our previous study^48^. We visually chose the obex (causal end point of quill-nib shaped calamus scriptorius, the inferior part of rhromboid fossa) as the anatomical reference for stereotaxic coordinates. The dorsomedial NTS, which is known as barosensory neurons gathers^49^, was targeted with regard to the reference (rostral: 0.3-0.6 mm; lateral: 0.4-0.8 mm; ventral: 0.3-0.8 mm) based on the widely known anatomy of rat brain^53^. Extracellular single-neuronal spikes were observed to monitor neural activity of the NTS with 16-channel silicone probe microelectrode (A1×16-Poly2s-5mm-50s-177-A16, NeuroNexus Technologies). Raw analog signals were converted into digital signals with sampling rate of about 24kHz (exactly 24414.0625 Hz) by a commercialized DAQ platform, which consists of a microprobe-headstage adaptor (ZCA-DIP16, TDT), a headstage (ZC16, TDT), and a neuro-digitizer (PZ5-32, TDT). The DAQ platform was protected from electromagnetic noise, especially power-source 60Hz noise and ground noise, by shielding our recording environment with a customized Faraday’s cage (copper-mesh and aluminum profiles). The electrical ground of DAQ system was connected to the cage and the DAQ output was transferred to a PC-controlled bioprocessor (RZ5, TDT) outside the cage through shielded optical cable. Real-time digital filters were applied on the neural data using a software suite (OpenEx, TDT): Highpass filter (>0.5 Hz) was applied to reject any long-term baseline fluctuations; Notch filter at 60 Hz was additionally applied to compensate the analog cancelling for electrical noise. The recorded neural activities were first save as TDT tank format and then transformed into MATLAB (Mathwork) data format ‘.mat’ by using TDT’s offline MatlabSDK.

### Haemodynamic recording

The real-time beat-to-beat arterial blood pressure was monitored through the intravenous catheter in femoral artery using a commercialized recording system (IX-RA-834, iWorx), which was administrated by its own software (LabScribe, iWorx), and a pressure transducer (BP-102, iWorx) placed at the level of subject heart. The catheter tubing was threaded out of the Faraday cage and connected to the transducer and this was to avoid for electrical noise from haemodynamic recording devices to interfere the neural recording. The blood pressure was recorded in 100 Sample/second and the event points such as stimulation on/off-set were manually marked on the time course. The transducer and catheter were filled with heparinized saline (heparin sodium: purchased from Merck(Sigma-Aldrich), product number: H5515-25KU; solution density: 100 IU/mL) to prevent blood coagulation that severely attenuates the pressure. Additionally, the blood diffused out into the catheter was perfused back into the artery using the solution about once an hour as well. The recorded raw pressure data was exported into the MATLAB data format ‘.mat’.

We calculated haemodynamic functions of the mean blood pressure, the heart rate, and the heart rate variability by post-processing the pressure recordings. The mean blood pressure (normally referred as the arterial BP in previous sections) was calculated by bandpass filtering the time course of raw pressure (high: >0.001 Hz and low: <0.5 Hz). The heart rate was calculated from the detected systolic peaks in highpass-filtered raw pressure signal (>1 Hz). We counted the number of beats in each time sampling step (0.01 s) as the instantaneous heart rate and elicited the heart rate by lowpass filtering the instantaneous heart rate with moving average filter (window: 3 s). As for the heart rate variability, we used the spectral power of heart rate. We calculated the spectrogram of heart rate for each 15 s time step with the Hanning window and 80% overlap among adjacent time steps. The low-frequency power of heart rate variability was the cumulative spectral power in the frequency range <0.15 Hz and the high-frequency power of heart rate variability in the frequency range 0.15-3 Hz (we empirically chose the ranges to clearly distinguish the difference responses between low and high frequency powers with considering various ranges reported in previous study^54-56^).

### Identification of single neurons

We post-processed the recorded NTS neural activities in offline to detect the single-neuronal spikes and appropriately cluster them to identify each single neuron (the process called ‘sorting’^57,58^). Before the sorting, we first removed stimulus-artifacts from the neural data with previously described ‘discard-and-treat’ approach^48,59^. We performed the sorting using an open source Klusta software suite^60^. The spikes were automatically detected from the filtered neural data (300-6000 Hz) and were clustered into putative single-neurons. The clusters were manually corrected if needed, using an open source Phy python package^61^. The toolkit provided the autocorrelogram of clusters that were calculated as the histogram of temporal lags around each spike (1 ms bin-step within 20 ms window range) and thus represented the refractory period of single neuron. We considered the cluster having peak values within the range within 2 ms as the capture of multi-unit activities rather than single-neuronal activity and excluded the clusters. Additionally, we excluded the clusters that did not have the V-shaped spike waveform or spread out the spikes at more than 10 channels, thereby considering as miss-detection of noise.

### Classification of NTS neurons

We divided the recorded and sorted NTS neurons into the responsive and non-responsive neurons to the solitary tract stimulation. The responsive neurons were chosen as showing stably identical responses to repeated stimulation trials. We evaluated the repeatability by using phase locking value calculated with firing rate of a neuron as 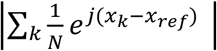 where N is the time length, *x*_*k*_ is the firing rate at time *k*, and *x*_ref_ is the reference signal at time *k*. For simplicity of calculation, we used a cosine wave as the reference signal that has a same period with the time course of repeated stimulation. Threshold of PLV >0.1 was empirically chosen to determine whether the neuron shows repeatable response. Interspike-interval (ISI) validated that the stimuli evoke the spikes. We presented the probability distribution of ISI and found peaks synchronized to the stimuli interval of 50 ms as shown in Supplementary Fig. 1b. We excluded non-responsive neurons and utilized only responsive neurons in further analyses. Moreover, we classified the responsive neurons into three subtypes with consideration of their heterogeneous responses to stimulation input; activated, deactivated, and complex. The activated neurons were ones showing the increased firing rate during stimulation, the deactivated neurons were ones showing the decreased firing rate during stimulation, and the complex neurons were ones showing both the increase and the decrease of firing rate during the stimulation.

### Inter-neuronal coupling analysis

To evaluate the joint activity induced by the interconnection among the NTS neurons, we calculated the pairwise cross-correlation and the shared variance^62^. The correlations were reported with the Pearson’s correlation coefficient by using MATLAB built-in function ‘corrcoef’ with inputs of firing rates of each neuron. Although the correlation coefficient elicited both positive and negative value due to heterogeneous responses of NTS neurons (see ‘Classification of NTS neurons’), we took the absolute value of correlation coefficients to only assess the strength of inter-neuronal coupling and prevent the mean across neurons being 0 that cannot be distinguished to the case of no coupling.

The shared variances were reported from the Factor analysis by using MATLAB built-in function ‘factoran’: **X** = **M** + **ΛF** + **E** where **X** is the k–by–n observed firing rate matrix (k: number of observations; n: number of neurons), **M** is the k–by–n observed mean matrix whereby column vectors are identical, **Λ** is the k–by–d factor loading matrix (d: number of common factors), **F** is the d–by–n common factor matrix, and **E** is the k–by–n independent specific factor matrix. The covariance matrix *cov*(**X** − **M**) can be expressed as: *cov* (**X** − **M**) = **LL**^**T**^ + *cov*(**E**). This elicits that the total variance is equal to the summation of the specific variance and the factor loading, and thus the shared variances of neuron ***i*** equal to the fraction of the factor loading relative to the total variance was calculated as:

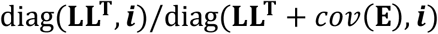

where diag(·, ***i***) is ***i***^th^ diagonal component of matrix. The shared variance stands for the fraction of variance of a neuron explained by the other neurons. Both the (absolute) correlation coefficient and the shared variance were compared to those of shuffled dummy data which were expected to have no entanglement. The shuffled dummies for neurons were generated by random permutation of time courses of firing rate.

### Dimensionality reduction for collective dynamics

The joint activities across single NTS neurons confined the degree of freedom in neuronal population dynamics and this characterized that the *n*-dimensional state space of neuronal population (*n* is the number of neurons) could be compressed into a low-dimensional subspace that effectively embeds the neural states for collective dynamics. The inherent noise into neural state representations in high-dimension makes it hard to understand the properties of collective dynamics^34,63^, and thus the redundant subspaces were removed by reducing the dimensions. We used a non-linear dimensionality reduction technique, called ‘Isomap’^64^, as previously described^32^ because it has been known to stably and effectively estimate the ‘neural trajectory’ in latent space compared to the linear approach such as principal component analysis. Before the dimensionality reduction, the firing rates of neurons were estimated by counting the number of spikes in the time step (0.5-0.7 s, empirically chosen for animals) and then filtering with the Gaussian kernel (window: 3 s; standard deviation: equal to time step). The firing rate patterns were normalized into the average value of maximums across stimulation trials. We performed the dimensionality reduction by using the ‘isomap’ class in Scikit-learn python package (neighbor number: 5). The Isomap used the geodesic distance matrix of the NTS neuronal neural state representations in high-dimensional coordinate, and then achieved the principal components of latent coordinate by calculating eigenvectors.

### Persistent homology

We used the persistent homology^32,34^ in topological data analysis to identify the geometric characteristics of collective representations lying on the latent space. This topological data analysis approach has been successfully used to revealing neural geometries such the sphere in primary visual cortex^65^, the ring of head direction cells^32^, and the toroid of grid cells^34^. Thus, we applied it onto discovering the topology embedded within the NTS. The Ripser python package^66^ was implemented to calculate the persistent homology. The calculation of persistent homology was as following. We considered the points of NTS neural activity in the dimension-reduced latent space as the 0-diameter balls in topological space. While increasing the scale, the geometrical structures that unify overlapped balls were detected. The essence of calculation was to analyse the geometrical ‘holes’ that are contained in the unifying structures at a given scale (for example, if a structure shapes ring, this implies there exists 1-dimensional hole). In formal notations, the persistent homology resulted in the properties of topological holes in Vietoris-Rips complexes^67^ (formal term for the unifying structure we use here). The properties of holes included the dimension and the scale range where holes persist. The dimension of holes was divided into three different homology groups^68,69^: 0-dimensional group H^0^(including no holes; ‘point’); 1-dimensional group H^1^ (including a hole made by single line; ‘ring’), and 2-dimensional group H^2^ (including a hallow space enclosed by single surface; including examples of ‘hallow sphere’ or ‘toroid’). The scale range is referred as ‘lifetime’. The longer lifetime implies the stable existence of hole, and thus confirms the topological characteristics. The Betti barcode graph visualized the lifetimes of holes and thus enabled to determine the geometrical shape of NTS neural trajectory.

### Effective dimension of latent space

Despite the latent space embedding by the dimensionality reduction (see ‘Dimensionality reduction for collective dynamics’), this providing of non-linear eigenvectors (principal components) of neural data did not specify the optimal dimension that captures the neural trajectory in latent space; this is because the maximum number of eigenvectors is theoretically equal to the high dimension of original coordinate. We achieved the effective dimension of NTS neural trajectory in the latent space by analysing the variance explained by each principal component and the correlation dimension of latent space. The explained variance along the order of principal components was computed as the percentage of variance of neural trajectory being projected on a principal component relative to the total variance of neural trajectory. The correlation dimension was calculated, as previously described^33,70^, by analysing the statistics of nearest neighbors in the neural data points. We acquired the geodesic distance matrix of latent space. The cumulative number of neighbors ‘*m*’ within a given distance threshold ‘*d*’ was counted while gradually increasing the distance threshold. The correlation integral was achieved as *C*(*d*) ≈ *m*/*N*^2^ where *N* is the total number of data. The correlation dimension was estimated with the slope of linear regression in the log-log graph between the correlation integral and the *d* such that *C*(*d*) = *kd*^*r*^ where *r* is the correlation dimension and *k* is a constant.

### Single-neuronal dependence on internal variables

We analysed the capture of single-neuronal activities by the external variable in physical domain (especially, the change in blood pressure) and the internal variable tracking along the neural trajectory. We used the internal variable defined as the ‘neural state phase’: the angle of point along the neural trajectory spanning on latent space, with adopting the polar coordinate system with the center point of trajectory as the reference origin. The capture of single-neuronal activities by the variables were quantitatively assessed with the fraction of variance explained by the tuning curve of neurons along the external and the interval variables and the mutual information between the single-neuronal activities and the variables. The tuning curves of single neurons along the external and the internal variable were calculated as the mean firing rate of neuron at a given value in 25 bins for each variable. The fraction of variance for each neuron was calculated as the ratio of variance for the changes in tuning curve relative to the total variance of single-neuronal activities. The mutual information for each neuron were calculated as previously described^71^. We analogously used this spatial information formula to compute the mutual information (MI) of a neuron for the external and the internal variables:

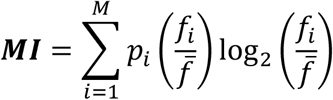

where *p*_*i*_ is the occupation ratio to be in *i*^th^ bin of variable, M is the total number of bins, *f*_*i*_ is the firing rate of the neuron at the *i*^th^ bin, and 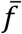 is the mean firing rate of the neuron for overall bins.

### Disentanglement of neuro-haemodynamic axis

We analysed the entanglement of input-driven haemodynamics into the latent space representations and compared it with the entanglement into the single-neuronal activities. To assess the entanglement, we performed the prediction of the input-driven haemodynamics with the neural data and used the prediction accuracy as the strength of entanglement. We used the linear regression for the prediction in both cases (with including a constant term as the bias) since decoding neuronal activities into the physical domain generally follows the linear relationships^38^. The regression was performed with a MATLAB built-in function ‘regress’ with 95% confidence interval. The measure of prediction accuracy was assessed using the goodness-of-fit of regression, specifically employing the coefficient of determination to quantitatively provide how successfully the neural data fit into the haemodynamics. The coefficient of determination of linear regression is equal to the square of Pearson’s correlation coefficient that has been widely used for the measure of accuracies in neuroscience and neural engineering applications such as modelling the stimulus-induced brainwave^59^ and decoding the behavior from cortex recording^72^. Even the coefficient of determination quantities the fraction of variance in ground-truth inputs explained by the prediction outputs, thereby representing the measure of robustness. Additionally, we quantified the error of prediction by calculating the squared errors between the measured haemodynamic perturbations and the regressed predictions.

For single-neuronal entanglement, each prediction accuracy of single neurons was calculated from a single-variable linear regression between the time course of firing rate and the haemodynamic perturbations.

As for the latent space representations, we used two-variable linear regression of the 1^st^ and 2^nd^ principal components into the haemodynamic perturbations (since we confined the effective dimension of latent space as 2D, as shown in Supplementary Fig. 2). This allowed to define ‘decoding vector’ consisting of the regression coefficients, and correspondingly ‘decoding space’, a 1D linear subspace (line subspace), consisting of the projections of collective neuronal neural state along the neural trajectory onto the decoding vector. The projection of the states onto the decoding vector designated the haemodynamic values. This could be expressed as following:

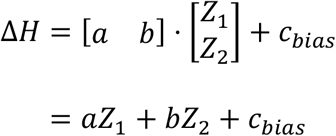

where Δ*H* is the change in haemodynamic functions such as blood pressure and heart rate, [*a b*]^**T**^ is the decoding vector (and the decoding space should be the *y* = *b/a x* line subspace in latent space), [*Z*_1_ *Z*_2_]^**T**^ is the neural state along the neural trajectory in 2D latent space, and *c*_*bibb*_ is a constant bias (The projection of neural state 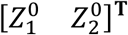 corresponding to Δ*H* = 0 onto the decoding vector is not the origin of decoding space, and this generally means that 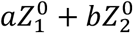 is not 0. Thus, we added the bias to adjust the origin).

### Normalization of neural trajectory

The inter-individually heterogeneous neural trajectories, as shown in Fig. 4a&b and Supplementary Fig. 4, were normalized by aligning the neural trajectories with combinations of linear transformation; ‘sliding’, ‘scaling’, ‘flipping’, and ‘rotating’ (Supplementary Fig. 4d) (thus, the combined transform was defined as a linear mapping function; *f*_*align*_: ℤ → 𝕐 where ℤ is original latent space spanning heterogeneous neural trajectories and 𝕐 is aligned(normalized) latent space).

The sliding adjusted the center points 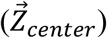 of ring-shaped neural trajectories into the origin point of latent space by subtracting the vector of center point from neural trajectory; 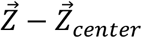. We estimated the center points as the mean of the maximum and the minimum values along each horizontal and vertical axis of latent space. The scaling weakly transformed the ring trajectories into the unit circle by normalizing each scale of the axes; 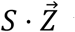 where 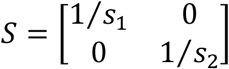, *s*_1_ is the horizontal scale, and *s*_2_ is the vertical scale. The scales of axes were estimated by the half of length between the maximum and the minimum values. These established a consistent geometrical reference frame of trajectories. The flipping and the rotating were to normalize the dynamical traces and the neuro-haemodynamic coupling protocols. The flipping unified the directions of input-driven temporal rotation along the trajectory into counter-clockwise, the direction of increasing the angle of neural state phase (see **‘**Single-neuronal dependence on external and internal variables’); 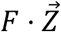 where 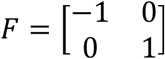. The rotating transformation involved in adjusting the decoding vector to be equal to the vertical axis; 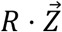 where 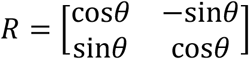 where 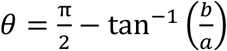 and 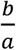 is the slope of decoding vector (see ‘Disentanglement of neuro-haemodynamic axis’). Following these, the normalized neural trajectory ([*Y*_1_ *Y*_2_]^**T**^ ∈ 𝕐) possessed the across-rat common protocol of neuro-haemodynamic coupling:

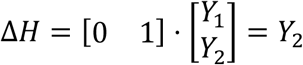

where [0 1]^**T**^ is the normalized decoding vector, and the values in vertical axis should be equal to the haemodynamic perturbations that were elicited as *aZ*_1_ + *bZ*_2_ + *C*_*bias*_ with unaligned and rat-specific neural trajectory. Additionally, the normalized coupling protocol exhibited that the normalized decoding vector is a unit vector of vertical axis and that the removal of bias. The sliding allowed the zero-point of ring-shaped neural trajectory that corresponds to no change in haemodynamics to be the origin point of latent space and this rejected bias; The scaling normalized the scale of neural trajectory along vertical axis in range from -1 to +1 and this matched in one-to-one relationship between the vertical value and the haemodynamic perturbations (which were normalized in the range from -1 to +1).

To assess the normalization across animal subjects, we measured the PLV and STD of polar distribution of neural state phase (*θ*). We made a polar distribution histogram that spanning in 30 bins, and compared the histogram between the unaligned and the aligned (normalized). The PLV and STD were measured for resting and recovery status that the neural state in the latent space should be static and stable. The PLV was calculated by 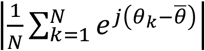 where *N* is total number of *θ* data, *θ*_*k*_ is the *k*^*th*^ *θ*, and 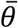 is the polar mean of *θ* distribution calculated by 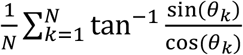, and the circular STD was calculated as 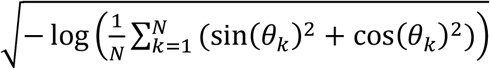.

### Modelling local neural circuit in NTS

To model the local neural circuit within the NTS, we designed the model structure with an artificial neural network, especially the RNN, which has been widely used to model the biophysically-plausible modelling of interconnected network in neuronal population^38,73^:

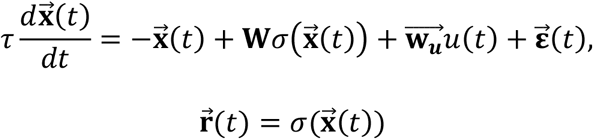

where 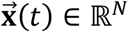 is the vector of membrane currents of *N* neurons (also called ‘activation’), 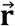 is the firing rates of neurons, *σ*(·) is the activation function of *tat h*(·), *τ* is the time constant of neuron, **W** is the *N*-by-*N* connectivity matrix, 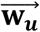 is the synaptic weight vector of stimuli inputs on neurons, *u*(*t*) is the time course of visceral sensation-mimicking stimulation, and 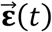 is the vector of independent Gaussian noise with variance of 1/*N*. We used the difference equation of the RNN model by discretizing with the time step Δ*t* equal to *τ* = 100 ms:

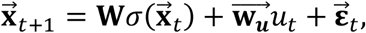

(and equivalently derived as 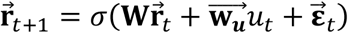 in the firing rate form^74^).

We embedded the latent representation into the RNN model as previously described^73^: 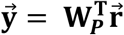 and 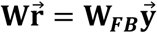 where 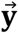 is the vector of neural state long the neural trajectory in the normalized latent space, **W**_***P***_ is the *N*-by-2 projection matrix from firing rate to the latent space, and **W**_***FB***_ is the *N*-by-2 recurrent feedback matrix that transforms the latent space into the synaptic recurrent inputs of neurons. This derived our final model equations:

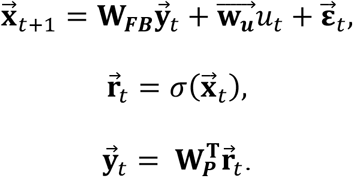

Before training the RNN, the vector 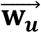 was pre-determined as the *N-*independent Gaussian distributions, the matrix **W**_***FB***_ was randomly generated with *N-*by-2 independent Gaussian distributions, and 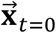 was initialized as 0. For the training, we optimized the *N*-by-2 matrix **W**_***P***_, which had the initial values of 0, using the recursive least square (RLS) appraoch^75^. The procedures of training started to calculate 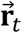 as 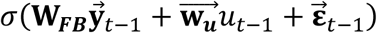, and then compared the estimation of neural trajectory 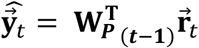 with the ground-truth neural trajectory 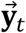. Finally, the 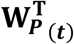 was updated using the estimation error 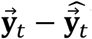^76^. The training data was the time course of 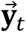 representing the neural trajectory in the 2D normalized latent space. We reformed the time course of single stimulation trial (basically, 20s resting time, 60s stimulation, and 20s recovery time; but could be changed the resting and recovery time according to the length of recordings in animal subjects) for the training data. The trial epoch was serially repeated at least 20 times and thus we made a long repeated epoch. This helped the precise fitting since the training algorithm progressed along the time axis. After training finished, we validated the fitted RNN by performing the feedforward simulation and verifying the accurate match between the simulated neural trajectory and the training data. The simulation was serially repeated 5 times and we verified the successful prediction for all repeats.

### H-BIND

To infer the viscerosensory input-driven haemodynamic perturbations, we used the NTS neural circuit model and the coupling protocol of neuro-haemodynamic axis. The NTS model calculated the input-driven collective dynamics within the NTS, and the dynamics was transformed into the haemodynamic representations using the coupling protocol. This procedure mathematically quantified the haemodynamics after visceral sensation, and thus the H-BIND identified the haemodynamic system using the neural computation mechanisms within the NTS.

The model architecture of H-BIND consisted of ‘encoder’, ‘NTS neural circuit model’, and ‘decoder’ (Supplementary Fig. 8a). The encoder transformed the changes in haemodynamic functions into the drift within the normalized latent space as known that the haemodynamic perturbations correspond to the neural trajectory along the unit circle in the normalized latent space. For the encoding, we first calculated the current phase (*θ*) of neural state in the latent space as sin^−1^ Δ*H* (where Δ*H* is the change in haemodynamic) to obtain the horizontal value of neural state as cos *θ*; its vertical value was equal to the Δ*H* as aforementioned (see ‘Normalization of neural trajectory’). Although sin^−1^ Δ*H* generated two possible values, only the *θ* was selected to satisfy monotonic increase since the neural trajectory only turns in counter-clockwise direction. The neural trajectory was obtained by calculating the temporal drift of neural state using the NTS neural circuit model. The model received both the current neural state and the stimulation as inputs, and then computed the new neural state in next time step. The neural trajectory was simply transformed back into the change in haemodynamics by the decoder since the vertical values of neural trajectory are equal to the haemodynamic perturbations. Thus, this allowed to predict the temporal dynamics of haemodynamics induced by stimulation.

The training of H-BIND identically means the training of NTS neural circuit model (since the encoding/decoding protocols are based on the determined neuro-haemodynamic coupling rules), and this required the neural trajectory as training data (see ‘Modelling local neural circuit in NTS’). However, the encoder could generate the neural trajectory from haemodynamics. This enabled to train the NTS neural circuit model with haemodynamic recordings (Supplementary Fig. 8b).

We assessed the performance of H-BIND to predict the input-driven haemodynamics using the k-fold cross-validation method^59^. We used the mean prediction accuracy across folds to indicate the H-BIND performance. The error of prediction was quantified with the distribution of squared error. Additionally, we assessed the prediction of neural trajectory by H-BIND in both aspects of radial and phasic accuracies.

### Data analysis and Statistics

Analyses were performed with custom scripts in MATLAB (R2022b) and Python (3.9.7) with Anaconda3 virtual environment (conda 23.1.0). We used open-source Python packages: numpy (1.21.5), scipy (1.7.3), ripser (0.6.1), scikit-learn (1.0.2), matplotlib (3.5.1), klusta (3.0.16), and phy (2.0b1). Python was used to detect and sort neuronal spikes and compute the latent space and its effective dimension. MATLAB was used for other analyses, including the preprocessing of electrophysiological neural and haemodynamics recording, the disentanglement of neuro-haemodynamic axis, the normalization of neural trajectory, and the construction of H-BIND.

Resources included all measured neurons that satisfied the classification criteria (see ‘classification of single neurons’) from 10 different rat subjects. No statistical standard was used to determine sample size for subjects and neruons. The experimental protocols of study did not require any experimental or control groups of animal, thereby not including random allocation and blinding tests. The statistical tests were all two-sided. This required the normality test with the Kolmogorov-Smirnov method. When rejecting its null hypothesis that the resources follow normal distribution, the Mann-Whitney U-test was performed; all cases we analysed rejected the null hypothesis. Two-sided resources, including parametric statistics, usually used boxplots with MATALB built-in function ‘boxplot’ with raw data scatters: cross-correlation and shared variance (measure vs. shuffle dummy as in Fig. 2b); fraction of variance and mutual information (external vs. internal variable as in Fig. 2f); prediction accuracy (single-neuron vs. neural trajectory as in Fig. 3f&g; unaligned vs. aligned as in Fig. 4d&f; empirical decoding vs. H-BIND as in Fig. 5d&e); if not the case, we specified so.

## Data availability

Data used in this project are available at https://osf.io/z7pj8/.

## Code availability

Custom scripts in MATLAB and Python are available at our GitHub repository (https://github.com/Ez0-9606/NTS_Neuro-haemodynamic)

## Acknowledgements

This research was supported by the Pioneer Research Center Program through the National Research Foundation of Korea funded by the Ministry of Science, ICT & Future Planning (2022M3C1A3081294); the National Research Foundation of Korea(NRF) grant funded by the Korea government(MSIT) (No. 2020R1A2C2005385).

## Author contributions

J.L. conceived and proposed the study. J.L. and J.M. designed and performed experiments. J.L. developed and performed analyses. J.L. and J.M. discussed and interpreted the results. J.L. and S.M.P. wrote and edited the manuscript. S.M.P. supervised the project. J.L., J.M., and S.M.P. obtained the funding.

## Additional information

Correspondence and requests for materials should be addressed to Sung-Min Park

