## Supplementary information for "Neural computation in the brainstem for visceral sensation-driven haemodynamics"

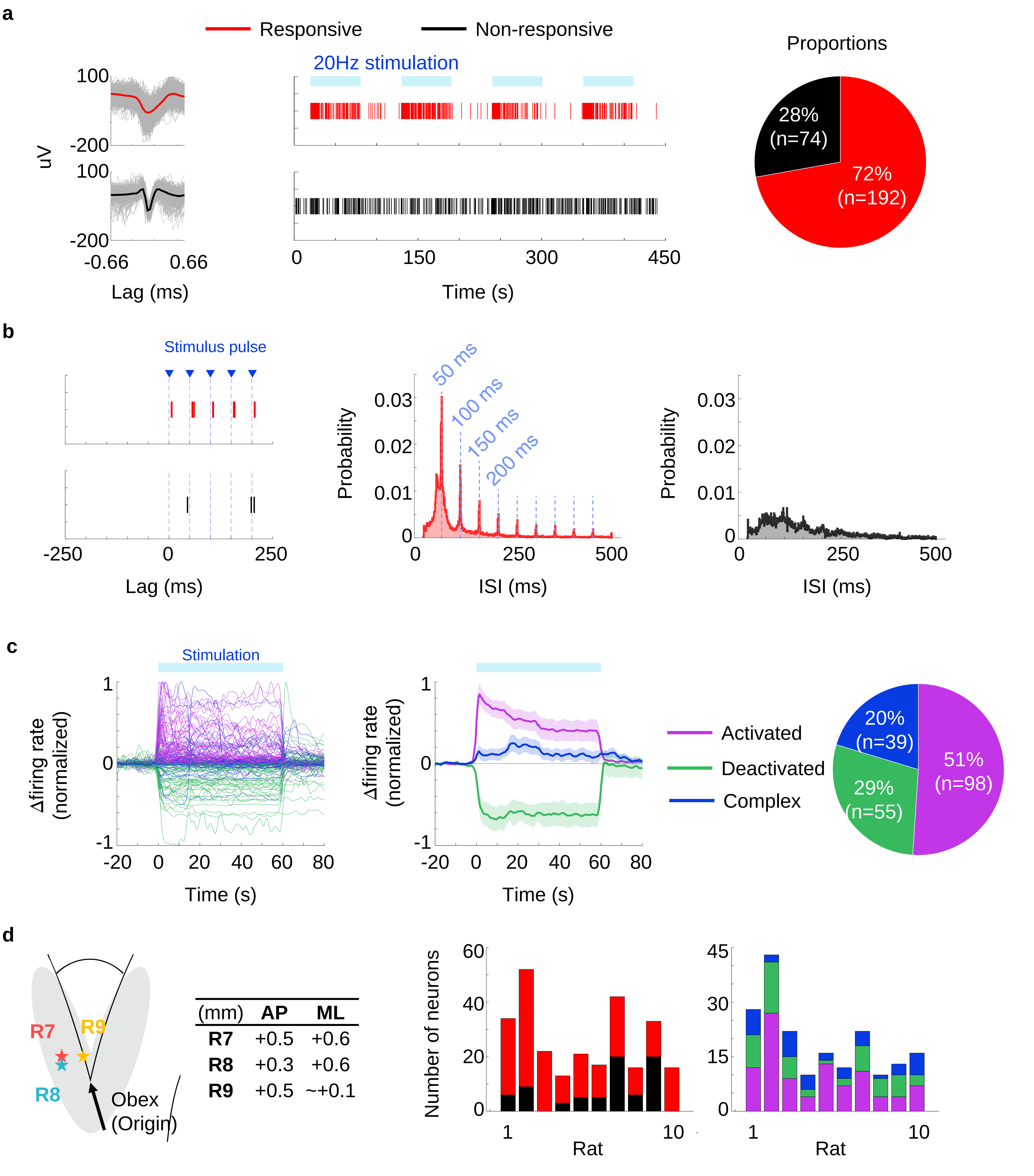


**Supplementary Fig. 1 | Properties and organization of NTS neuronal population.**

**Supplementary Fig. 1 | Properties and organization of NTS neuronal population.**

**a**, Recording of single-neuronal spike activities responding to barosensory-mimicking external stimulation. Representative illustration of spike waveforms of responsive and non-responsive neurons (left), raster plot (middle). The overall proportion observation from 10 rats (right). **b**, Different inter-spike interval (ISI) of responsive and non-responsive neurons. The responsive neuron presents stimuli-induced activation compared to the non-responsive neurons. **c,** Configuration of responsive neurons. The observe responsive neurons were sub-divided by the firing rate patterns into three groups of ‘activated’, ‘deactivated’, and ‘complex’. Right diagram presents the proportion of sub-groups in total 192 responsive neurons. **d,** Inter-animal heterogeneity of NTS measures. Representative recording location difference of three rats (left). Rat-specific proportion of responsive and non-responsive neurons (middle) and configuration of responsive neurons (right).

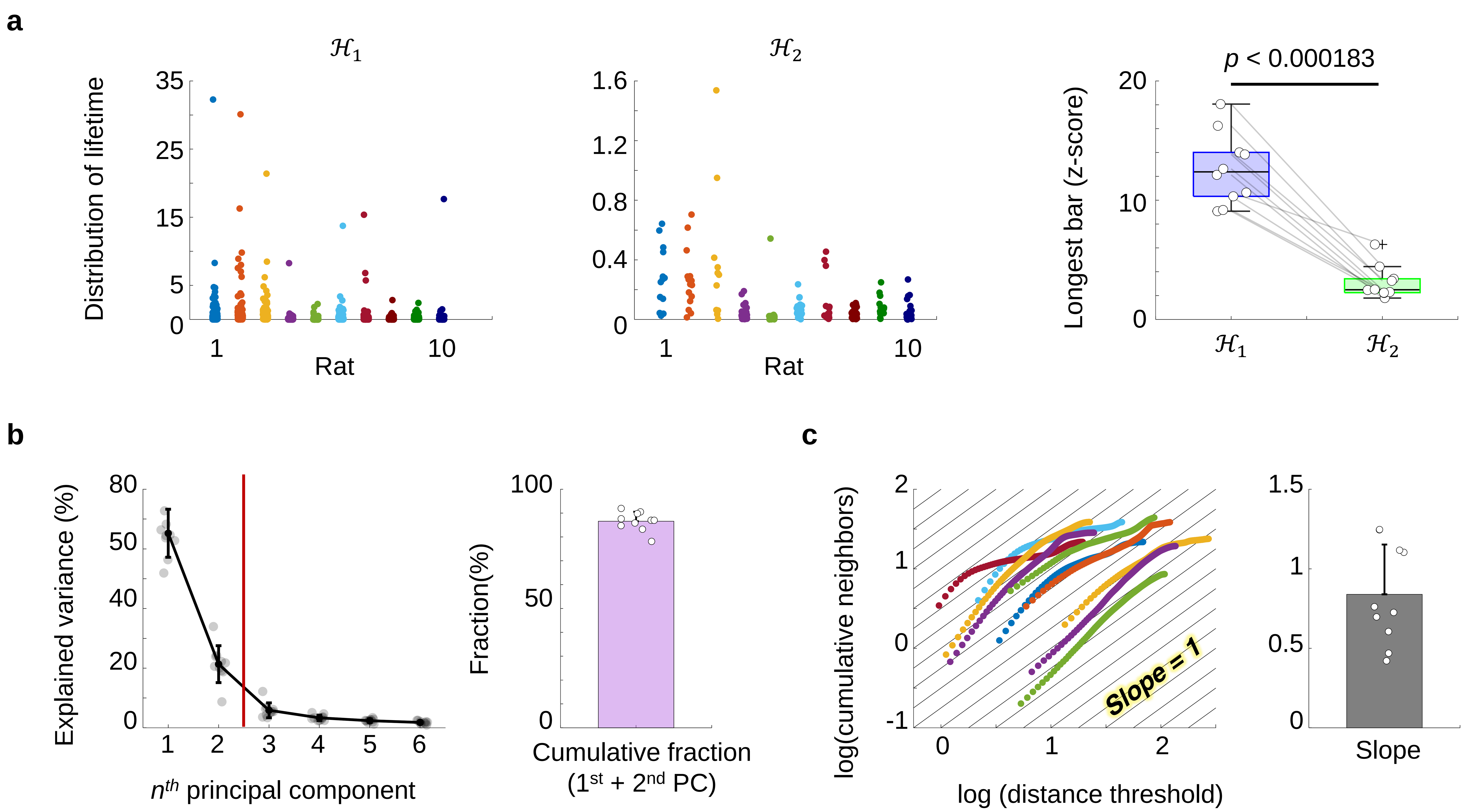


**Supplementary Fig. 2 | Effective dimension of latent space for neural trajectory from the NTS.**

**Supplementary Fig. 2 | Effective dimension of latent space for neural trajectory from the NTS**.

**a**, Rat-specific distribution of lifetime of Betti barcode in the persistence homology group $H_1$ and $H_2$ (*n* = 10 rats; box: median ± IQR for normalized (z-score) lifetime of the longest bars). **b**, Capture of explained variance by the principal component (PC) of latent space (*n* = 10 rats; bar: mean ± standard deviation (STD) for the cumulative explained variance with $1^{st}$ and $2^{nd}$ PC). **c**, Correlation dimension which was evaluated by the slope of linear regression between radius versus number of neighbors graph (*n* = 10 rats; bar: mean ± standard deviation (STD) for the slope).

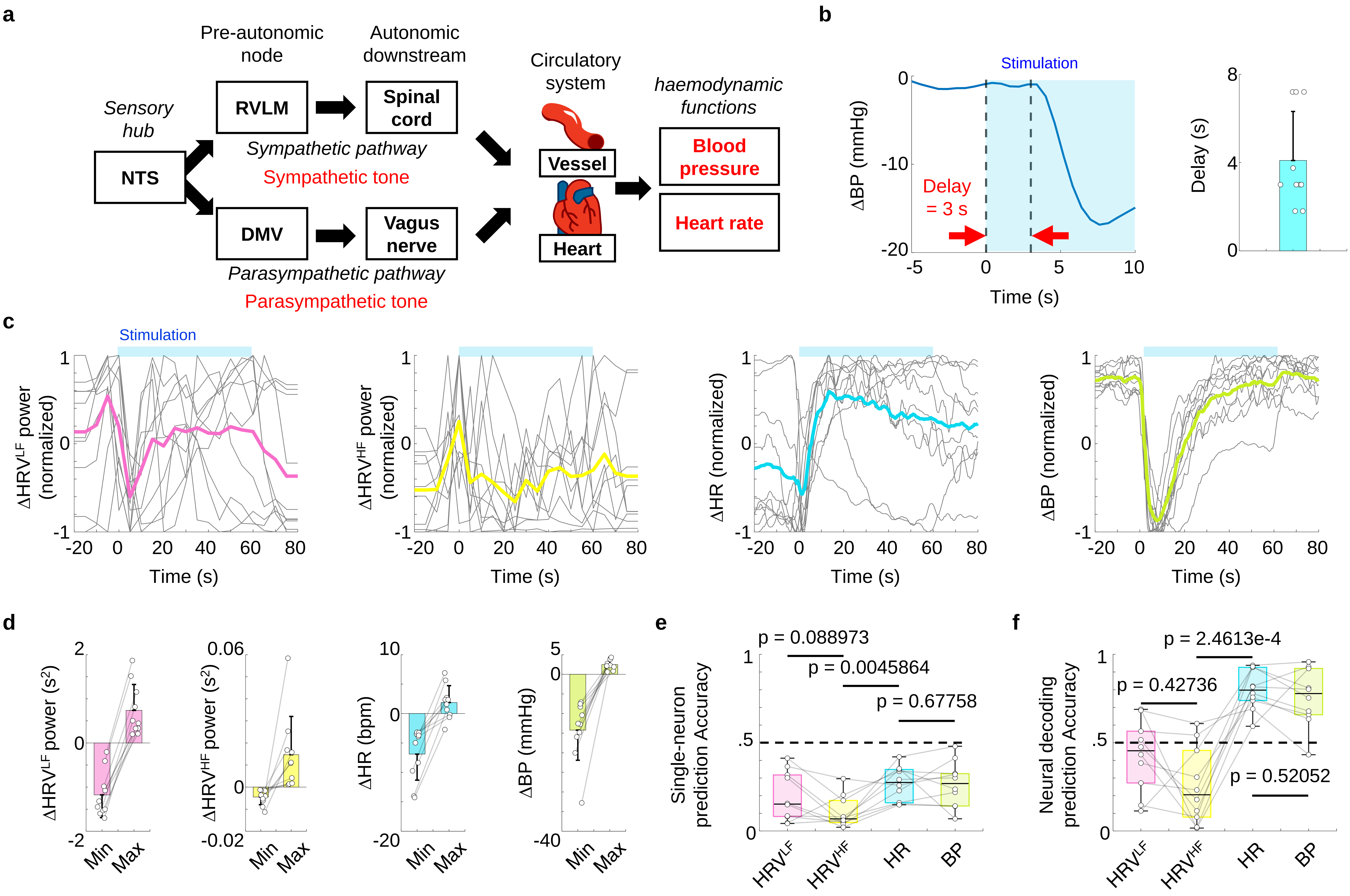


**Supplementary Fig. 3 | Coupling between various haemodynamic functions with the NTS.**

**Supplementary Fig. 3 | Coupling between various haemodynamic functions with the NTS.**

**a**, Illustration of neurological anatomy between the NTS and peripheral haemodynamic functions. From the NTS, sympathetic pathway passes through rostral ventrolateral medulla (RVLM) and parasympathetic pathway passes through dorsal motor nucleus of vagus (DMV), and then autonomic tones modulate circulatory organs resulting haemodynamic perturbations. **b-f,** Statistical analyses for solitary tract stimulation-induced haemodynamic perturbations (*n* = 10 rats). **b**, Time delay until the point of blood pressure (BP) decreasing after the NTS receiving stimulation (bar: mean ± standard deviation). **c**, Normalized time courses of various haemodynamic functions; low- and high-frequency powers of heart rate variability (HRV), heart rate (HR), and BP (power of heart rate variability stands for the autonomic tones: low- and high-frequency for each of sympathetic and parasympathetic tome). **d**, Maximum and minimum changes of haemodynamic functions (box: median ± interquartile range (IQR) for each of maximum and minimum). **e-f**, Prediction of haemodynamic functions by single-neuronal activities (**e**) and by neural decoding with neural trajectory(**f**) (box: median ± IQR).

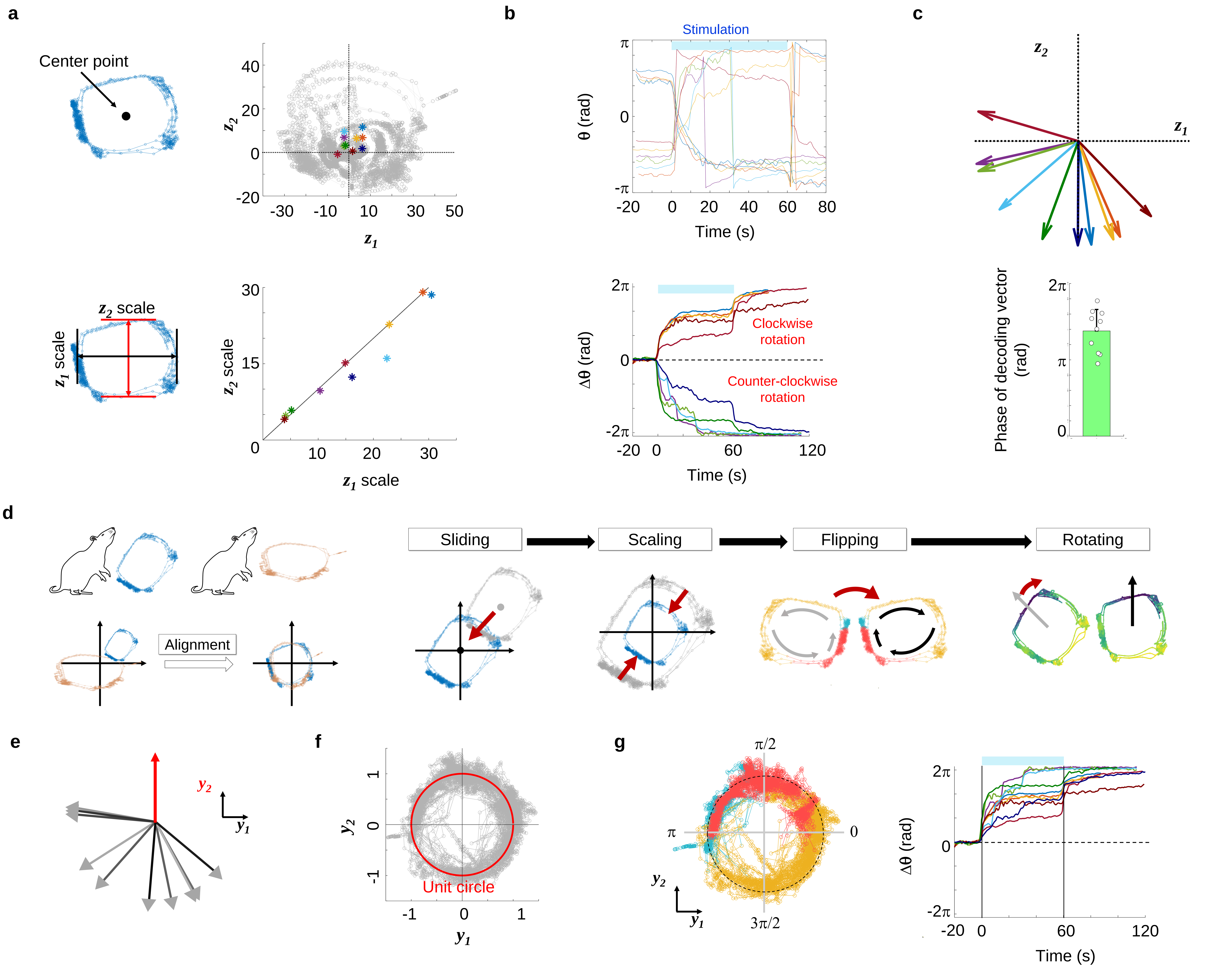


**Supplementary Fig. 4 | Normalization of heterogeneous neural trajectory across rats.**

**Supplementary Fig. 4 | Normalization of heterogeneous neural trajectory across rats.**

**a-d**, Heterogeneity in rat-specific neural trajectories (*n* = 10 rats). **a**, Geometric heterogeneity shown by the location of centor points and scales ($\boldsymbol{z_1}$ and $\boldsymbol{z_2}$ are the 1st and 2nd principal component of latent space). **b**, Heterogeneity in dynamics of neural state along neural trajectory indicated by neural state phase ($\theta$). **c**, Heterogeneity in haemodynamic coupling with by the direction of decoding space. **d**, Across-rat normalization procedure of neural trajectory through alignment. **e**, Normalization of decoding vector to be equal to vertical axis of aligned latent space ($\boldsymbol{y_1}$ and $\boldsymbol{y_2}$). **f**, Geometrical normalization that projects neural trajectories onto the unit circle in the aligned latent space. **g**, Dynamical normalization of neural trajectory shown by the distribution of $\theta$ (cyan: pre-stimulation; peach: post-stimulation; yellow: stimulation). (Right) Time courses of $\Delta\theta$ in identical rotational direction after normalization.

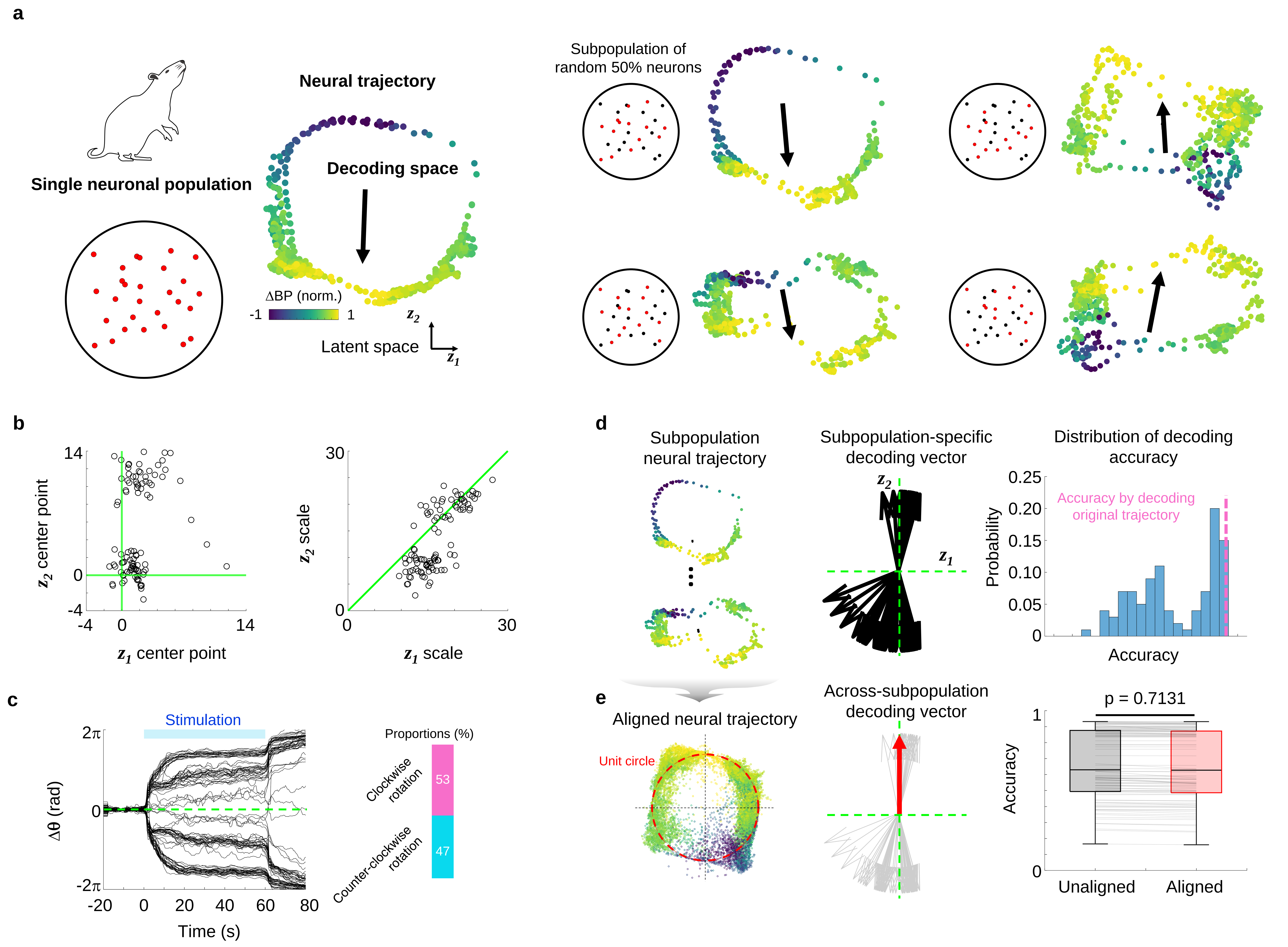


**Supplementary Fig. 5 | Distortion of neural trajectory due to sampling subpopulation.**

**Supplementary Fig. 5 | Distortion of neural trajectory due to sampling subpopulation.**

**a,** Various distortions of original neural trajectory by sampling 50% random subpopulation from original population (14/28 neurons from Rat 1; and haemodynamic coupling to change in blood pressure (ΔBP)). Right representatives show various neural trajectories according to subpopulations; one almost identical to original but another severely distorted. **b-d**, Distortion of subpopulation neural trajectory (*n* = 100 random subpopulations). **b**, Distortion in geometrical properties ($\boldsymbol{z_1}$ and $\boldsymbol{z_2}$ are $1^{st}$ and $2^{nd}$ principal component). **c**, Distortion in dynamical properties shown by time courses of change in neural state phase (Δθ). **d**, Distortion in haemodynamic couplings. **e**, Normalization of various subpopulation trajectories (left: topology aligned on unit circle; middle: decoding vector aligned on vertical axis; right: DBP prediction accuracies of neural decoding with aligned neural trajectories compared with those of neural decoding with unaligned trajectories (*n* = 100; box: median ± interquartile range).

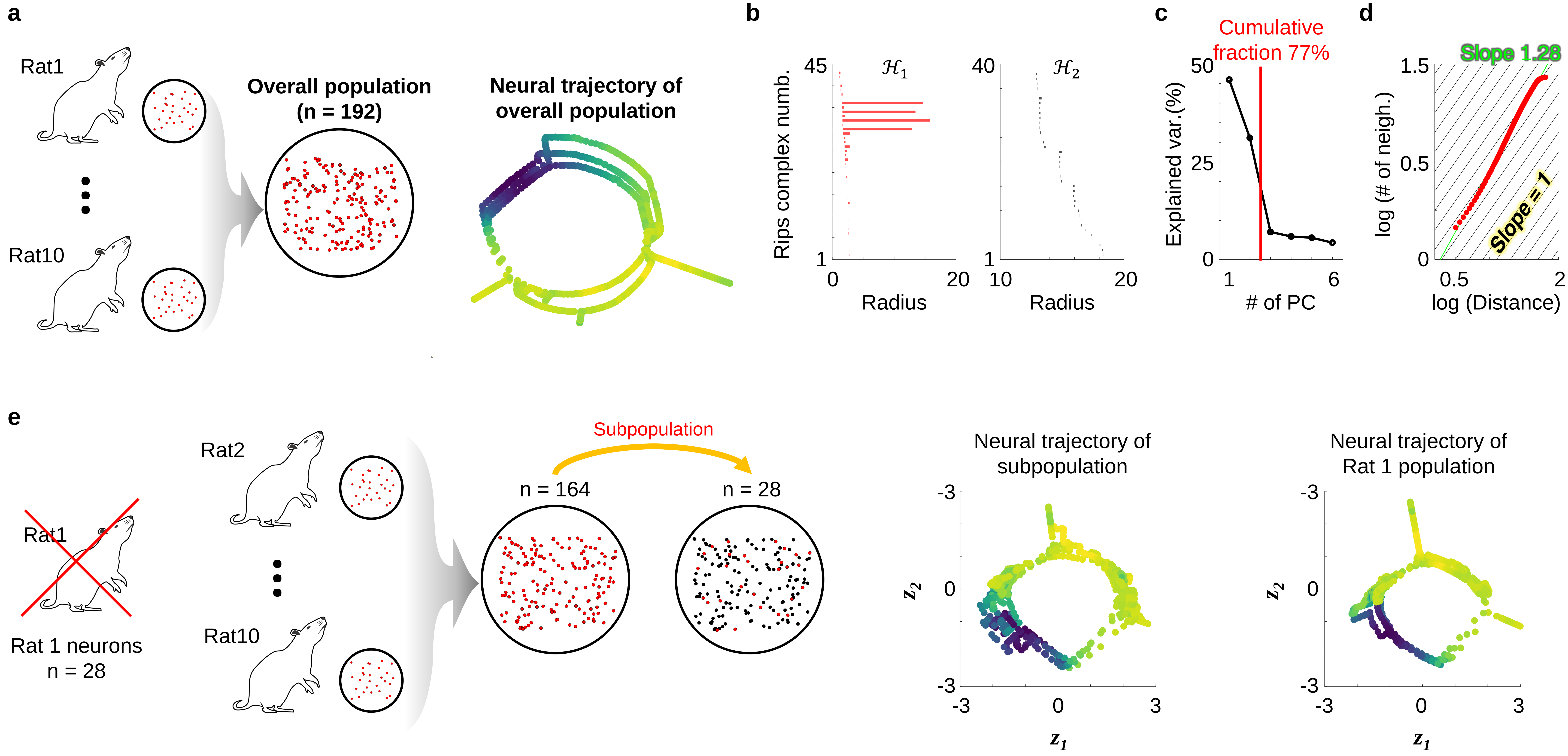


**Supplementary Fig. 6 | Commonality of NTS neuronal population across rats.**

**Supplementary Fig. 6 | Commonality of NTS neuronal population across rats**.

**a**, Neural trajectory of the across-rat overall neuronal population which gathers all measured neurons in 10 rats. **b-d**, Effective dimension analyses for the overall neural trajectory (similarly to Supplementary Fig. 2). **b**, Persistent homology. **c,** Explained variance by principal components. **d**, Correlation dimension. **e**, Almost accurate reconstruction of neural trajectory of Rat 1 by the subpopulation sampled from the neuronal population gathering the other rats except for Rat 1.

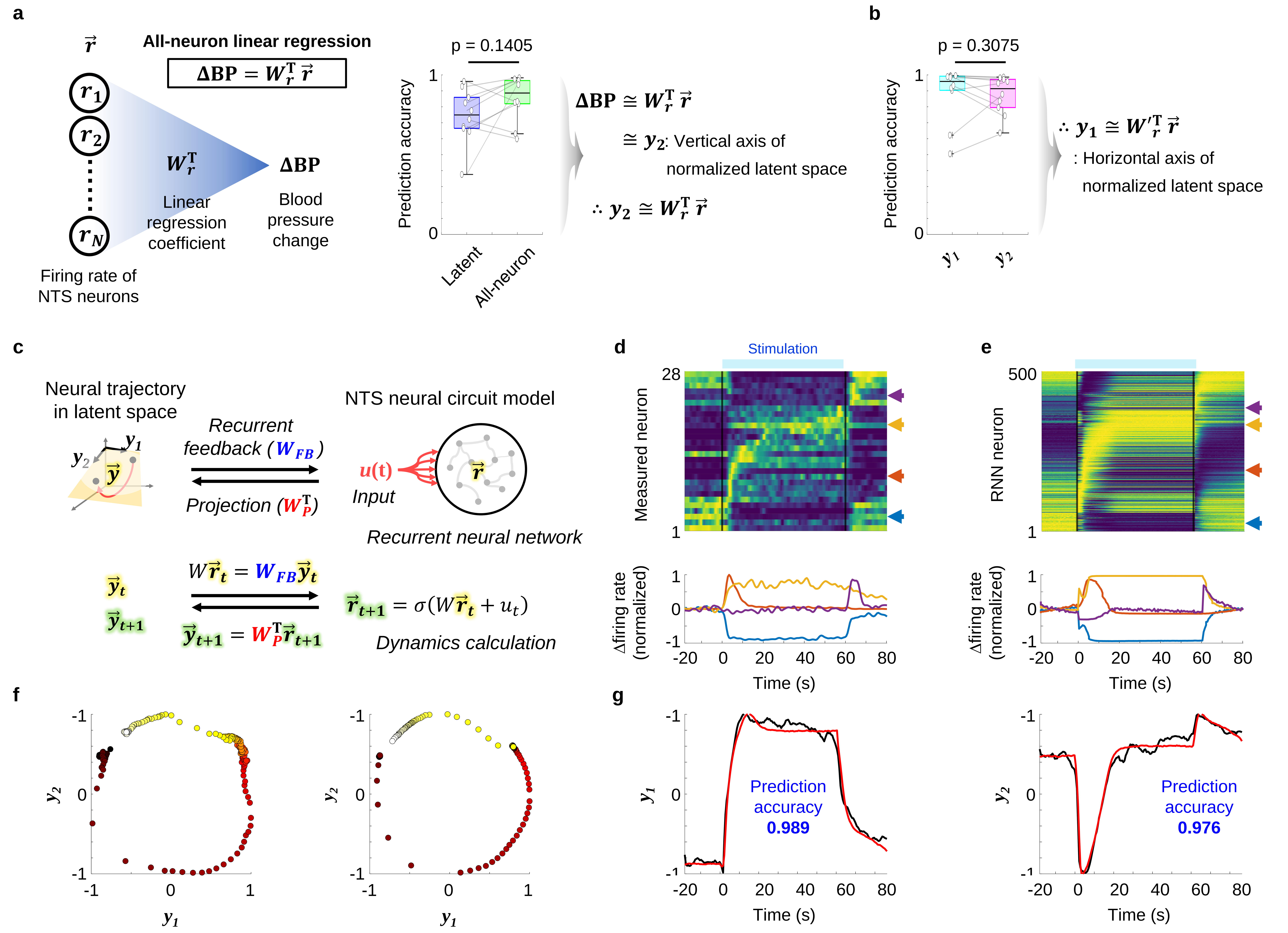


**Supplementary Fig. 7 | Modelling NTS neural circuit using neural trajectory measures.**

**Supplementary Fig. 7 | Modelling NTS neural circuit using neural trajectory measures**.

**a-b,** Feasibility of approximation of dynamics in normalized latent space dynamics using linear regression of single-neuronal activities. **a**, Prediction of change in blood pressure (ΔBP) using the linear regression of single-neuronal activities did not differ with that of the neural decoding using normalized latent space, indicating that the changes in vertical axis of normalized latent space can be exhibited by linear regression of single-neuronal activities. **b**, Accuracy of linear regression of single-neuronal activities to predict the change in horizontal axis in the normalized latent space did not significantly differ to the accuracy of linear regression for the changes in vertical axis. **c**. Model of NTS neural circuit to embed neural trajectory in (normalized) latent space ($\vec{y}$) into single-neuronal activities ($\vec{r}$), using the findings in **a** and **b**; $\boldsymbol{W_{FB}}$: feedback matrix from latent space to single-neuronal activities, $\boldsymbol{W_P}$: projection matrix from single-neuronal activities to latent space, $\boldsymbol{s}$: activation function 'tanh', and $\boldsymbol{W}$: synaptic weight matrix compromised as $\boldsymbol{W_{FB}}$ $\boldsymbol{W_P}^{\mathrm{T}}$. **d-f.** Comparison between NTS neural circuit model and the measured NTS neurons. **d**, Measured single-neuronal activities. **e**, Simulated single-neuronal activities with NTS neural circuit model. **f-g**, Comparison of measured neural trajectory with the simulation by NTS neural circuit model. **f**, Presentation in latent space. **g**, Time courses in 1st ($\boldsymbol{y_1}$) and 2nd ($\boldsymbol{y_2}$) principal component in normalized latent space.

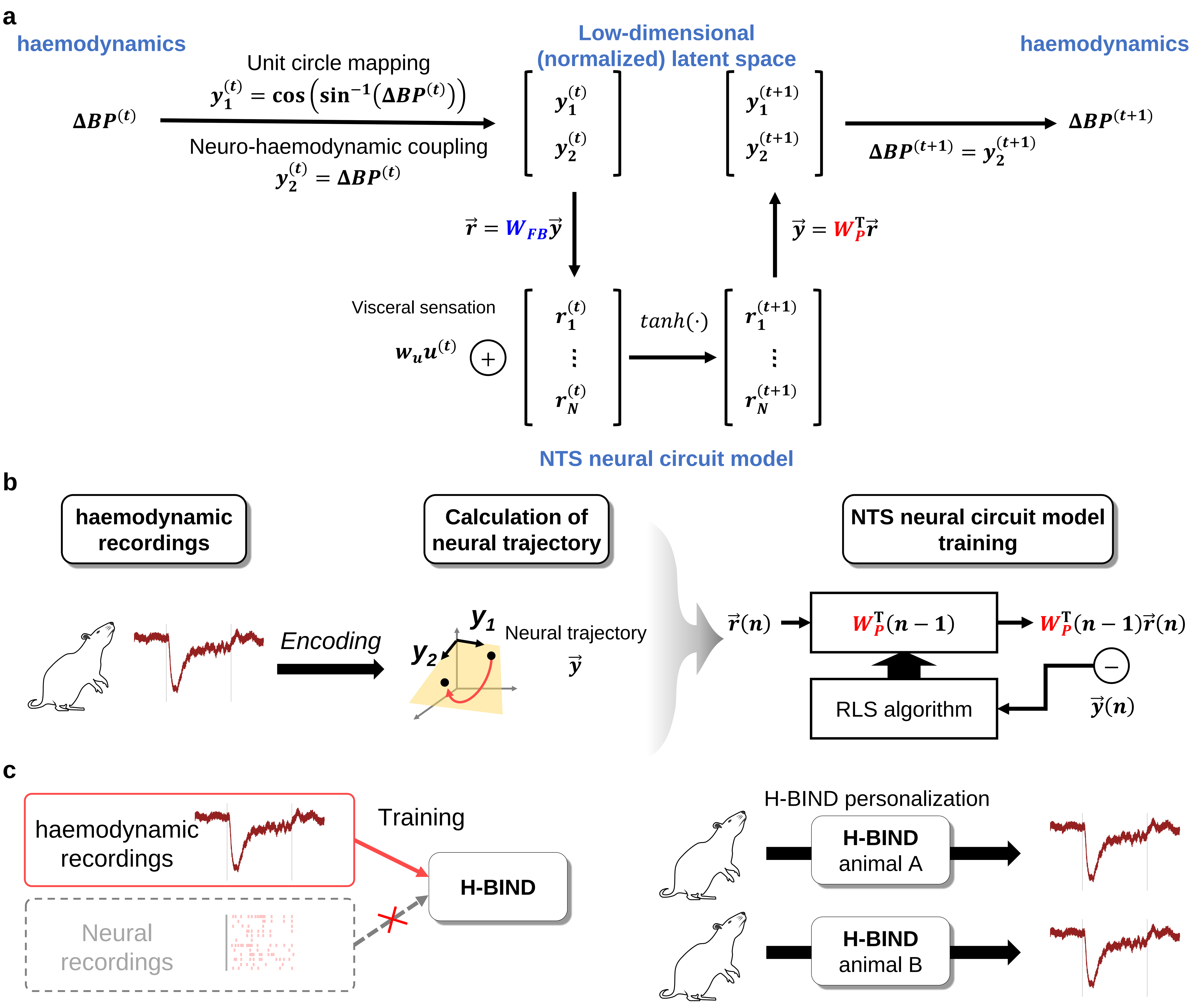


**Supplementary Fig. 8 | Model architecture, construction procedures, and advantages of H-BIND.**

**Supplementary Fig. 8 | Model architecture, construction procedures, and advantages of H-BIND**.

**a**, Overall model architecture of haemodynamics model based on barosensory input-driven neural dynamics (H-BIND). **b**, Training of recurrent neural network (RNN) that stands for NTS neural circuit model in H-BIND. **c**, Illustrations to explain advantages of H-BIND. Left shows that the training of H-BIND needs only haemodynamic recordings and right shows that the H-BIND can be rat-specifically optimized.

a

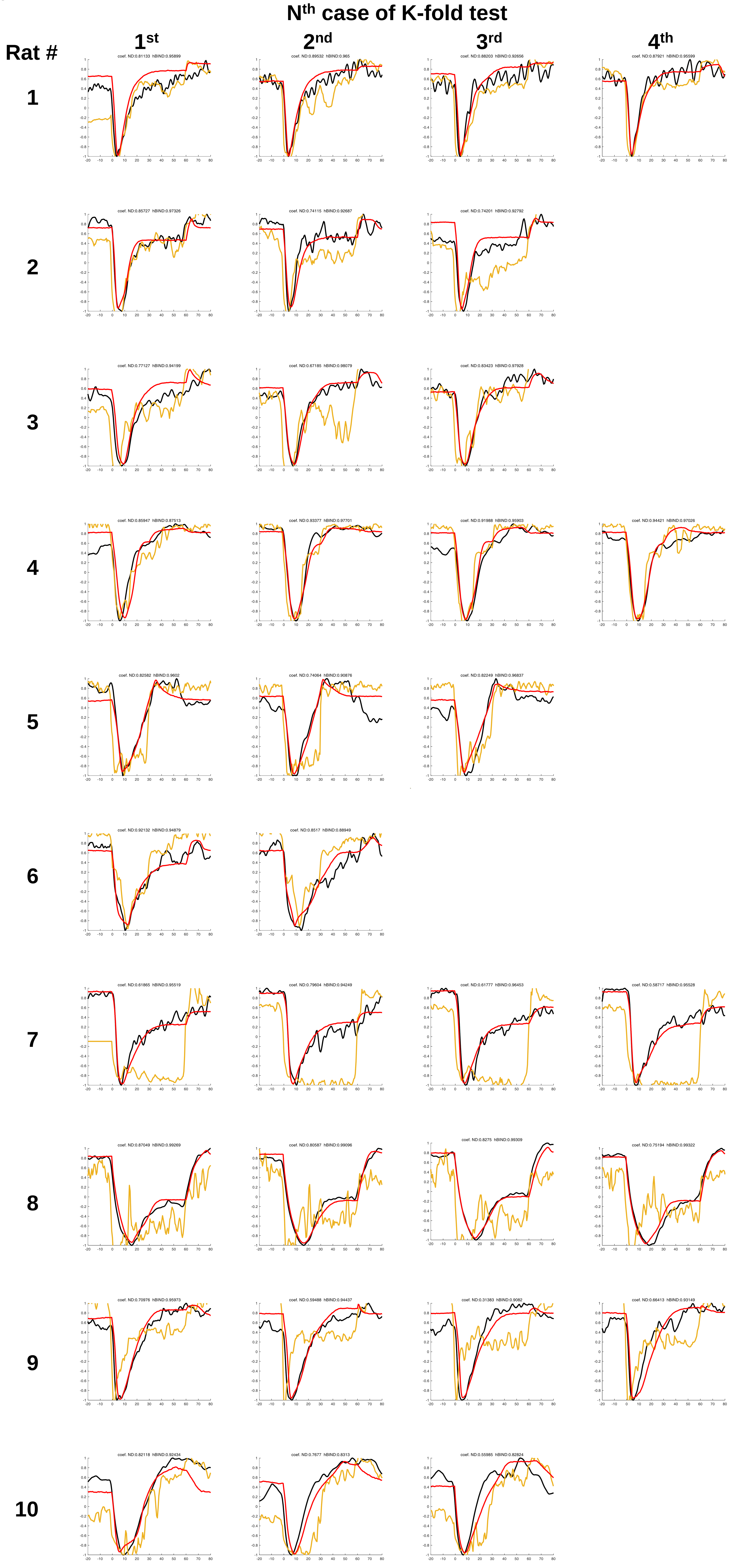


b

| rat | 1st fold | | 2nd fold | | 3rd fold | | 4th fold | |
|---|---|---|---|---|---|---|---|---|
| 1 | 0.811 | 0.959 | 0.895 | 0.965 | 0.882 | 0.927 | 0.879 | 0.956 |
| 2 | 0.857 | 0.973 | 0.741 | 0.927 | 0.742 | 0.928 | | |
| 3 | 0.771 | 0.942 | 0.672 | 0.981 | 0.834 | 0.979 | | |
| 4 | 0.859 | 0.875 | 0.934 | 0.977 | 0.920 | 0.959 | 0.944 | 0.970 |
| 5 | 0.836 | 0.960 | 0.741 | 0.909 | 0.822 | 0.968 | | |
| 6 | 0.921 | 0.949 | 0.852 | 0.889 | | | | |
| 7 | 0.619 | 0.955 | 0.796 | 0.942 | 0.618 | 0.965 | 0.587 | 0.955 |
| 8 | 0.870 | 0.993 | 0.806 | 0.991 | 0.828 | 0.993 | 0.752 | 0.993 |
| 9 | 0.710 | 0.960 | 0.595 | 0.944 | 0.314 | 0.908 | 0.664 | 0.931 |
| 10 | 0.821 | 0.924 | 0.768 | 0.831 | 0.560 | 0.828 | | |

**Supplementary Fig. 9 | Performance of H-BIND to predict change in blood pressure compared to the empirical decoding with the measured NTS neural trajectory.**

**Supplementary Fig. 9 | Performance of H-BIND to predict change in blood pressure compared to the empirical decoding with the measured NTS neural trajectory**.

**a**, Time courses of prediction for change in blood pressure haemodynamics model based on barosensory input-driven neural dynamics (H-BIND) for each $N^{th}$ case k-fold test (for 10 rats), compared to the measured, and the prediction by empirical neural decoding. **b**, Comparison of prediction accuracy between H-BIND and the empirical neural decoding for change in blood pressure.
